# Efficient Inference on a Network of Spiking Neurons using Deep Learning

**DOI:** 10.1101/2024.01.26.577077

**Authors:** Nina Baldy, Martin Breyton, Marmaduke M. Woodman, Viktor K. Jirsa, Meysam Hashemi

## Abstract

The process of making inference on networks of spiking neurons is crucial to decipher the underlying mechanisms of neural computation. Mean-field theory simplifies the interactions between neurons to produce macroscopic network behavior, facilitating the study of information processing and computation within the brain. In this study, we perform inference on a mean-field model of spiking neurons to gain insight into likely parameter values, uniqueness and degeneracies, and also to explore how well the statistical relationship between parameters is maintained by traversing across scales. We benchmark against state-of-the-art optimization and Bayesian estimation algorithms to identify their strengths and weaknesses in our analysis. We show that when confronted with dynamical noise or in the case of missing data in the presence of bistability, generating probability distributions using deep neural density estimators outperforms other algorithms, such as adaptive Monte Carlo sampling. However, this class of deep generative models may result in an overestimation of uncertainty and correlation between parameters. Nevertheless, this issue can be improved by incorporating time-delay embedding. Moreover, we show that training deep Neural ODEs on spiking neurons enables the inference of system dynamics from microscopic states. In summary, this work demonstrates the enhanced accuracy and efficiency of inference on networks of spiking neurons when deep learning is harnessed to solve inverse problems in neural computation.

## 1. Introduction

Neural computation involves the complex information processing performed by spiking neurons, where their interactions within neural circuits contribute to higher-level computations and cognitive functions. Consequently, the cornerstone of theoretical neuroscience lies in the use of mathematical models to understand the intricacies of neural computation (Dayan and Abbott, 2005; Hertz, 2018). This approach enables researchers to explore the complexities of neural processes, thereby revealing insights into the mechanisms that drive brain function and behavior. These mathematical models can range from simple idealized representations of single neurons (Hopfield, 1982; Izhikevich, 2003) to complex network models that simulate the interactions within neural circuits (Marder, 1998; Sussillo, 2014; O’Leary et al., 2015; Bittner et al., 2021). The biological neural computation forms the basis for the brain’s ability to execute various inference tasks, such as recognizing patterns, making decisions, and generating responses to stimuli. This capability relies on the collective behavior of spiking neurons, where the interactions and coordination among these neurons within neural circuits enable the brain to process, integrate, and interpret sensory information, ultimately leading to higher-level cognitive functions and adaptive behavior (Kandel et al., 2000; Gerstner et al., 2014; Friston and Kiebel, 2009; Friston et al., 2017).

Mean-Field (MF) models serve as effective computational abstractions that represent the collective behavior of large populations of neurons, while maintaining a degree of mathematical tractability (Amari, 1977; Wilson and Cowan, 1973; Jirsa and Haken, 1996; David and Friston, 2003; Deco et al., 2008; Hutt et al., 2015; Coombes and Byrne, 2018; Bandyopadhyay et al., 2021; Cook et al., 2022). Hence, MF models facilitate the study of information processing and computation within the brain, which underlie cognitive processes such as perception, memory, and learning. Nevertheless, deriving and validating MF models from spiking neural networks present many challenges, particularly when assessing how the interaction between individual neurons and parameters leads to macroscopic behavior that aligns with the averaged activity of neural populations.

Recently, an analytically-driven MF model of spiking neurons has been formulated, which can effectively describe all potential macroscopic dynamical states of the network, including states of synchronous spiking activity (Montbrió et al., 2015). However, the operation of such complex systems (governed by coupled nonlinear differential equations) is determined by the selection of (biological or phenomenological) parameters, which when set in a specific configuration, give rise to a measurable signature of a computation (Achard and De Schutter, 2006; Sussillo, 2014). Analyzing MF models and comparing their emergent dynamics against a network of neurons involves solving inverse problems to ascertain the optimal parameter setting. This process requires swift and robust outcomes to inform real-time decisions and also to deal with observation and dynamical noise. Yet, even in the simplest models, there can be a degenerate relationship between the model parameters and its overall emergent function (Edelman and Gally, 2001; Prinz et al., 2004; Alonso and Marder, 2019), making the inverse problem more challenging.

Maintaining the inter-dependency between parameters by traversing across scales adds an additional layer of complexity to the validation process. It is crucial to distinguish between genuine, biologically relevant correlations, and artificial correlations that may arise from the inference process or modeling assumptions. Furthermore, due to the intricate nature of computation within neural circuits, it becomes intractable to analytically derive MF models that include more biological realism (such as adaption, neuromodulation, extra-synaptic transmission, and E/I ratios). Statistical inference offers an efficient and adaptable approach to solving the inverse problem by identifying approximate parameter distributions that are responsible for generating computations in a biologically realistic model (Achard and De Schutter, 2006; Liepe et al., 2014; Lueckmann et al., 2017; Gonçalves et al., 2020; Bittner et al., 2021; Młynarski et al., 2021).

The performance of statistical inference algorithms depends on the task, and there is no universally best algorithm for different inverse problems. Therefore, we conducted a benchmarking analysis against state-of-the-art optimization and Bayesian estimation algorithms to discern their respective advantages and limitations. In practice, optimization methods are commonly used to quickly determine unknown quantities through a single point estimate (Mendes and Kell, 1998; Nocedal and Wright, 1999; Kelley, 1999; Floudas and Gounaris, 2009). These methods involve iteratively adjusting parameters to minimize or maximize an objective function, scoring the model’s performance against observed data (e.g., through minimizing distance errors or maximizing correlation; Banga and Balsa-Canto (2008); Tashkova et al. (2011); Svensson et al. (2012); Hashemi et al. (2018)).

The Bayesian approach offers a principled method for making inferences, predictions, establishing relationships between parameters, and quantifying uncertainty in the decision-making process (Gelman et al., 1995; Bishop, 2006; Gelman et al., 2020; van de Schoot et al., 2021). In Bayesian modeling, all model parameters are treated as random variables and their values are subject to variation based on their underlying probability distributions. Such probabilistic techniques provide the full posterior distribution of unknown quantities hidden in the underlying data generating process. The uncertainty and inter-dependency in Bayesian estimation are naturally quantified by assigning a probability distribution to each parameter (known as the prior distribution), which is then updated based on the information provided by the data (referred to as the likelihood function). To conduct a fully Bayesian procedure, the state-of-the-art MCMC method is adaptive Hamiltonian Monte Carlo (HMC; Duane et al. (1987); Neal (2010); Hoffman and Gelman (2014)), which utilizes gradient information to avoid random walk behavior. This enables efficient sampling from high-dimensional distributions that may exhibit strong correlations (Betancourt, 2017).

Simulation-Based Inference (SBI; Cranmer et al. (2020); Brehmer (2021)) or likelihood-free inference (Papamakarios and Murray, 2016; Brehmer et al., 2020) leverages deep generative models to conduct approximate Bayesian estimation, using low-dimensional data features that are generated by random simulations (Gonçalves et al., 2020; Lueckmann et al., 2021; Boelts et al., 2022). In this efficient approach, a simple base probability distribution (prior) is transformed into a more complex distribution (posterior), through a sequence of invertible transformations (i.e., Normalizing Flows; Rezende and Mohamed (2015); Papamakarios et al. (2019a)). Notably, it allows for direct estimation of joint posterior distributions, bypassing the need for MCMC sampling (Greenberg et al., 2019; Papamakarios et al., 2019b). Moreover, expressive deep generative models have the potential to capture parameter nonlinear relationships between parameters and multi-modalities in the distributions (Hashemi et al., 2023).

Data-driven methods for learning dynamical models from time-series data have been extensively researched for several decades (Juang, 1994; Ljung, 1998; Brunton et al., 2016; Linderman et al., 2017; Duncker et al., 2019; Koppe et al., 2019; Sip et al., 2023). Instead of relying on discretized maps, Neural Ordinary Differential Equations (Neural ODEs; Chen et al. (2018)) form a new family of deep neural network models for modeling continuous-time dynamics. Neural ODEs define the vector fields and ODE solution as a black-box differential equation solver, allowing for uncovering the dynamics of a system even when the governing equations are unknown (Dupont et al., 2019; Biloš et al., 2021). This data-driven approach involves parameterizing system dynamics as continuous functions, enabling smooth and uninterrupted modeling of temporal evolution (Yan et al., 2019; Kim et al., 2021). Neural ODEs naturally adapt to varying time intervals and can accommodate fluctuations in the frequency of data observations (Zhu et al., 2022; Goyal and Benner, 2023).

Through an exploration of the aforementioned methods, we demonstrate that global optimization algorithms, such as Differential Evolution algorithm, offer fast and accurate point estimation of the true generative parameters when the dynamical noise is absent. However, when dealing with dynamic evolution that are subject to noise, SBI using deep neural density estimators emerges as the superior approach, outperforming other algorithms, such as adaptive HMC sampling. Additionally, when dealing with missing data (such as population firing rate) in state-space modeling, HMC fails to capture the dynamics of bistable switching behaviour. Instead, SBI is able to accurately recover the diverse dynamics in the phase-space representation. Nevertheless, this approach may lead to an overestimation of uncertainty and correlation between parameters, which can be mitigated by using time-delay embedding technique to improve the results. We validate this by employing a MF model of quadratic integrate-and-fire (QIF) neurons, which demonstrates that the interdependencies between parameters are maintained when traversing across scales. Finally, we demonstrate that training deep Neural ODEs on spiking neurons enables the inference of vector fields at macroscopic level. This allows for the prediction of emergent behaviors and system dynamics based on the microscopic state of the spiking neurons.

## 2. Materials and methods

### 2.1. A mean-field model of spiking neurons

The quadratic integrate-and-fire (QIF) neurons are a class of simplified computational models that are extensively used to study the dynamics of spiking neurons (Gerstner and Kistler, 2002; Izhikevich, 2007). In the QIF model, the membrane potential of neuron evolves according to a quadratic differential equation until it reaches a threshold, at which point the neuron emits a spike and the potential is reset.

Montbrió et al. (2015) have proposed a MF model that accurately describes macroscopic states of populations of firing neurons. This mechanistic model derives the firing rate equations for networks of heterogeneous, all-to-all coupled QIF neurons, which is exact in the thermodynamic limit, i.e. for large numbers of neurons. Specifically, when considering specific distributions of heterogeneity, the Lorentzian ansatz yields a nonlinear system of two ordinary differential equations for the firing rate *r* and mean membrane potential *v* of the neuronal population:

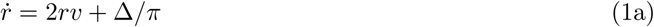

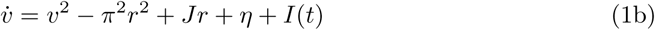

where *η* is the average excitability, *J* denotes the synaptic weight, and Δ indicates the spread of the neuronal excitability distribution in the neural population. Depending on the parameter settings, the phase diagram exhibits three qualitatively distinct regions: a single stable node, which represents a low-activity state; a single stable focus (spiral), which generally corresponds to a high-activity state; and a region of bistability, where both low and high firing rates can coexist (see Figure S1). This model has succeeded in establishing an exact correspondence between the time evolution of firing rate of the network and the underlying microscopic state of the spiking neurons (Montbrió et al., 2015).

### 2.2. Inference methods

We validate the mean-field approximation by comparing it against detailed spiking neurons, using various parameter estimation inference methods. To evaluate these methods, we use synthetic data generated from the MF model and a network of QIF neurons. Inference methods investigated in this work can be broadly divided into two classes:

(i) Optimization methods that return a point estimate of best fit based on the minimizing of a cost function, such as chi-squared error criterion defined by

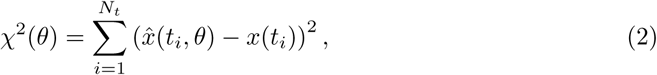

where *x*(*t_i_*) denotes the observed data at time points *t_i_* with *i* ∈ {1, 2, … , *N_t_*}, and *x̂*(*t_i_*, *θ*) represents the corresponding model prediction. Here *θ* ∈ {*η, J,* Δ} is the set of unknown parameters, and the set of observation combines activity of both *r*, *v* (unless it is missing). Assuming no prior information and a Gaussian likelihood function with uncorrelated noise, this casts as a Maximum Likelihood Estimation (MLE) problem (Hashemi et al., 2018).
(ii) Bayesian methods return a posterior distribution of parameters, *p*(*θ* | *x*), which represents an ensemble of parameter sets that are plausible given the observed data. Given the data *x* and model parameters *θ*, Bayes rule defines the posterior distribution as

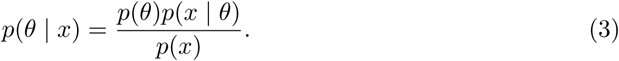

The prior information *p*(*θ*) is typically determined before seeing the data (through beliefs and previous evidence). The likelihood function *p*(*x* | *θ*) represents the probability of some observed outcomes given a certain set of parameters (the information provided by the observed data). The denominator *p*(*x*) = *∫ p*(*x* | *θ*)*p*(*θ*)*dθ* represents the model evidence or marginal likelihood, which amounts to simply a normalization factor.

From the optimization methods, we use the global search algorithms that incorporate a bio-inspired random search principle: Differential Evolution (DE; Storn and Price (1997); Price (1999)), and Particle Swarm Optimization (PSO; Kennedy and Eberhart (1995); Eberhart and Kennedy (1995)). These algorithms do not require an initial guess for the parameters or the gradient information of the objective function. We also consider Bayesian Optimization (BO; Snoek et al. (2012); Shahriari et al. (2015)), an algorithm that constructs a probabilistic model for the objective function using Gaussian processes. This approach allows for the integration of uncertainty in the optimization process.

From the Bayesian methods, we compare the results of two state-of-the-art Bayesian computation algorithms: Hamiltonian Monte Carlo (HMC; Duane et al. (1987); Neal (2010)) which is unbiased and exact in infinite runs, and a Simulation-Based Inference (SBI; Cranmer et al. (2020); Brehmer (2021)) which approximates the posterior using deep generative models. Here, generative modeling is an unsupervised machine learning method to model a probability distribution based on the samples drawn from that distribution. From the Bayesian methods, we also report the results of Maximum a Posteriori (MAP) estimation.

### 2.3. Hamiltonian Monte Carlo

Markov Chain Monte Carlo (MCMC) is a powerful class of computational algorithms used for sampling from a distribution, in which the sampling process does not require knowledge of the entire distribution, making it a versatile tool. (Andrieu et al., 2003; Murphy, 2022; McElreath, 2020). MCMC is unbiased and asymptotically exact, in the limit of infinite runs. Hamiltonian Monte Carlo (HMC; Duane et al. (1987); Neal (2010)) is a gradient-based MCMC designed to avoid random walk behaviour, and it can efficiently sample from high-dimensional distributions that may exhibit strong correlations (Betancourt, 2017). However, the efficiency of HMC is sensitive to the algorithm parameters.

In this study we use a self-tuning variant of HMC (known as the No-U-Turn Sampler; Hoffman and Gelman (2014)) from a high-level statistical modeling tool called Stan (Carpenter et al., 2017). In particular, the NUTS calibrates the number of steps and step-size of the leapfrog integrator (in solving the Hamiltonian equations of motion) during a warm-up phase to achieve a target Metropolis acceptance rate. For more details see Betancourt (2013); Baldy et al. (2023). Moreover, Stan offers alternative methods such as MAP estimation using L-BFGS optimization, automatic differentiation for efficient gradient computation, and various diagnostics to assess the convergence of the inference process (see https://mc-stan.org).

### 2.4. Simulation-Based Inference

Simulation-Based Inference (SBI) conducts efficient Bayesian inference for complex models when the calculation of the likelihood function is either analytically or computationally intractable (Cranmer et al., 2020; Brehmer, 2021). In computational models, where the data can be generated through stochastic simulations, SBI leverages repeated simulations from the generative model and employs probabilistic machine learning to estimate a target probability distribution. Instead of directly sampling from distributions using MCMC or explicitly evaluating the likelihood function, SBI overcomes these challenges by using deep neural density estimators, such as Masked Autoregressive Flows (MAF; Papamakarios et al. (2017)). These density estimators learn an invertible transformation between the distributions of parameters and low-dimensional data features at a very low computational cost, to efficiently sample from distributions.

Taking the prior distribution *p*(*θ*) over the parameters *θ*, a limited number of *N* simulations are generated for training step from the generative model i.e., 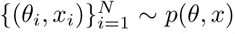. After the training step, we are able to quickly estimate the approximated posterior *q_ϕ_*(*θ* | *x*) with learnable parameters *ϕ*, so that for the observed data *x_obs_*: *q_ϕ_*(*θ* | *x_obs_*) *≃ p*(*θ* | *x_obs_*). For more details see Gonçalves et al. (2020); Hashemi et al. (2023).

The methods for SBI often include a sequential training procedure, which adaptively guides simulations to yield more informative estimates (Papamakarios et al., 2019b; Lueckmann et al., 2019; Durkan et al., 2020; Wiqvist et al., 2021; Deistler et al., 2022). In particular, Sequential Neural Posterior Estimation (SNPE; Greenberg et al. (2019); Gonçalves et al. (2020)) dynamically refines the proposals, network weights, and posterior estimates to learn the relationship between model parameters and the observed summary statistics of the data. In this study, we used SNPE with a single round to take advantage of an amortized strategy; After incurring an initial computational cost for the simulation and training steps to learn all the joint posterior distributions, then the posterior can be quickly estimated from any new observations (by a forward pass through neural networks) without any additional computational overhead or further simulations.

### 2.5. Time-delay embedding

Time-delay embedding is a commonly used technique for characterizing dynamical systems based on limited measurements, time-series analysis, and prediction (Takens, 1981). In time-delay embedding, the reconstruction of a latent high-dimensional system relies on incorporating incomplete measurements along with a temporal history of preceding measurements to create a comprehensive representation (Kennel et al., 1992; Hirsh et al., 2021). In a subsequent analysis, we challenge the inference process by assuming that the firing rate activity *r* is not directly observed (missing data problem). Instead, to improve the inference, we recovered the latent time-series *r_rec_* from the observed activity *v*, which is coupled to *r* according to Eq. 1. By expanding on a method introduced by Abarbanel et al. (1994), which primarily focuses on predicting physical variables in time-delay embedding, we removed the assumption that we have the access to training data points from the hidden time-series. Instead, we leverage our understanding of the generator to simulate data pairs (*r, v*) that can be used for training purposes.

Time-delay embedding requires the setting of hyperparameters, such as the delay (time lag), *T*, and the dimension of the embedding space, *d*. These hyperparameters are typically set prior to training, often based on the minimization of mutual information to determine the appropriate delay and the false nearest neighbors method to determine the number of embedding dimensions (Kennel et al., 1992; Tan et al., 2023). However, in the present application, we have found that the hyperparameters suggested by these methods were not the most effective in achieving an accurate fit. Instead, we select hyperparameters that minimize the mean square error of the fit to simulated data, with *T* = 160 points (i.e., 0.16 *sec*) and *d* = 12. To do this, a set of 100 pairs of coupled time-series (*r, v*) was simulated, with an observation noise intensity of 0.1 and varying parameters (Δ, *η, J*). These pairs were then used to infer regression coefficients that closely match *r* to the delay embedding space representation of *v*.

### 2.6. Neural ODEs

Neural ODEs are a set of machine learning techniques that allow to reconstruct the phase-space of a dynamical system from a training set of observations (Chen et al., 2018). In a first analysis, we trained a Neural ODE on time-series from the MF model, and then applied the same method to the data generated by a network of QIF spiking neurons. If **x**(*t*) is the vector of state variables governed by the dynamical system **ẋ** = *f* (**x**, *θ*, *I_ext_*), one can use a Neural ODE to approximate the function *f* with an artificial neural network *F_ϕ_*, yielding the corresponding dynamical equation 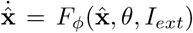. *F_ϕ_* is learned through backpropagation, minimizing the loss function for each training example of length *T* :

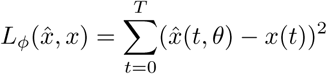

where *x̂* and *x* are times-series generated by *F_ϕ_* and *f*, respectively, with the same integration scheme (e.g., Heun’s method). In this study, a Neural ODE was implemented in JAX (Bradbury et al., 2018) by constructing a multi-layer-perceptron (MLP) with one hidden layer of 16 units, and with hyperbolic tangeant activation functions. The cost function was the average loss functions across training examples 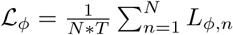. At each training iteration, the initial (*r, v*) values of each segment is fed to the Neural ODEs and the forecast trajectory according to current state of the *F_ϕ_* enters the cost function, then minimized through backpropagation using the Adam optimizer.

### 2.7. Software and algorithmic setup

We use Python implementations of the optimization and Bayesian algorithms: DE from SciPy (Virtanen et al., 2020), PSO from the toolkit PySwarms (Miranda, 2018), and BO from the package Bayesian Optimization (Nogueira et al., 2014). DE is set with population size of 10, and the maximum iterations of 500. PSO is set with 10 particles, and 500 iterations. BO is set with 50 random initialisation, 500 iterations, and kappa=100 (higher values increase exploration).

The MAP estimation is obtained using a quasi-Newton L-BFGS optimizer from Python interface to Stan (Carpenter et al., 2017), an open-source probabilistic programming language for Bayesian modeling and statistical inference. We also use Stan’s implementation of the NUTS for fully Bayesian estimation by HMC, with Metropolis acceptance rate of 0.8, maximum tree depth of 10, warm-up iterations of 1000, and 500 sampling phase iterations.

Finally, for SBI we used SNPE from the PyTorch-based SBI toolkit (Tejero-Cantero et al., 2020). SNPE was run for a single round with 100,000 simulations, unless specified otherwise. For training step, we used a MAF, with 5 autoregressive layers, each with two hidden layers of 50 units. The set of data features includes the statistical moments of time-series up to the fourth order (mean, standard deviation, skewness, and kurtosis), and the peak properties (number and location of first peak, as shown in Figure S2).

Parameter bounds for optimization, and uniform prior for Bayesian inference, were set as: Δ ∈ [0.1, 5], *η* ∈ [*−*10, *−*3], *J* ∈ [5, 20]. For the simulation of Eq. 1, we used Euler-Maruyama integration for *t* = 100 *sec* with a time step of *dt* = 1 *msec*. The ground truth parameters were set as *η* = *−*4.6, *J* = 14.5, and Δ = 0.7, ensuring that the system is in a bistable regime, unless specified otherwise. Moreover, a step current with amplitude of 3 *v* from time 30 *sec* to 60 *sec* was added to the *v* variable. Each simulation took around 0.01 *sec* to run using a Just-In-Time (JIT) compiler. To generate stochastic dynamics, a zero-mean Gaussian noise with *σ* = 0.1 was added to the both *r*, and *v* variables. The prior on dynamical noise was set as *σ* ∈ [0, 1]. For comparison with the MF model, we ran a network of 10^4^ all-to-all connected QIF neurons using the Brian simulator (Stimberg et al., 2019).

The model simulation and parameter estimation were performed on a Linux machine with 3.60 GHz Intel Core i7-7700 and 8 GB of memory.

## 3. Results

We present the results of the optimization and Bayesian algorithms for three cases: (i) inferring from exact deterministic synthetic data, (ii) inferring from stochastic synthetic data driven by dynamical noise, and (iii) inferring with stochastic synthetic data when the activity for *r* is unknown (missing data). We report and compare the goodness-of-fit using the root mean square error (RMSE) to the true (observed or hidden) time-series and parameters, the variance in the estimation process, and the computational cost.

Before running inference process, we conduct a sensitivity analysis to assess the structural identifiability of the parameters. This analysis involves calculating local sensitivity coefficients, which measure the effect of small adjustments in a parameter on the model’s output while keeping all other parameters constant. A small change in a sensitive parameter leads to a significant alteration in the model output, indicating its identifiability. Conversely, when there are no alterations in the model output despite adjustments in a parameter, it suggests the parameter’s non-identifiability. To assess the identifiability of the parameters, we used the profile likelihood (Raue et al., 2009; Wieland et al., 2021) and Hessian matrix, a metric describing the local curvature of a function based on its second partial derivatives (Hashemi et al., 2018, 2023). Our results indicate that all three parameters Δ, *η*, and *J* can be identifiable, however, the profile likelihood obtained using Eq. 2 reveals the presence of numerous local minima (see Figure S3). This highlights the difficulty in attaining the global minimum during the inference process.

### 3.1. Inference on deterministic data

Here, we compare the inference results of different algorithms when we have complete observations of both state variables (*r, v*), and the system operates without any noise (see Figure 1). The observed time-series and trajectories in phase-plane, which exhibit a bistability between a stable node and a stable focus, are illustrated in Figure 1**A**. From the results demonstrated in Figure 1**B**, it can be seen that all the algorithms qualitatively reproduce the bistability behaviour in the phase-plane. However, when evaluating their performance by estimating the true generative parameters, all optimization methods, except DE, get stuck in a local minima (Figure 1**C**). The results indicate that DE, SBI and HMC algorithms correctly recover the ground-truth parameters, the former as an almost exact point estimate (Figure 1**C**, top panels), while the latter two, SBI and HMC, yield full posterior distributions (Figure 1**C**, bottom panels). In terms of uncertainty quantification, HMC offers a slightly more informative posterior distribution compared to SBI (see Figure S4).

**Figure 1.**
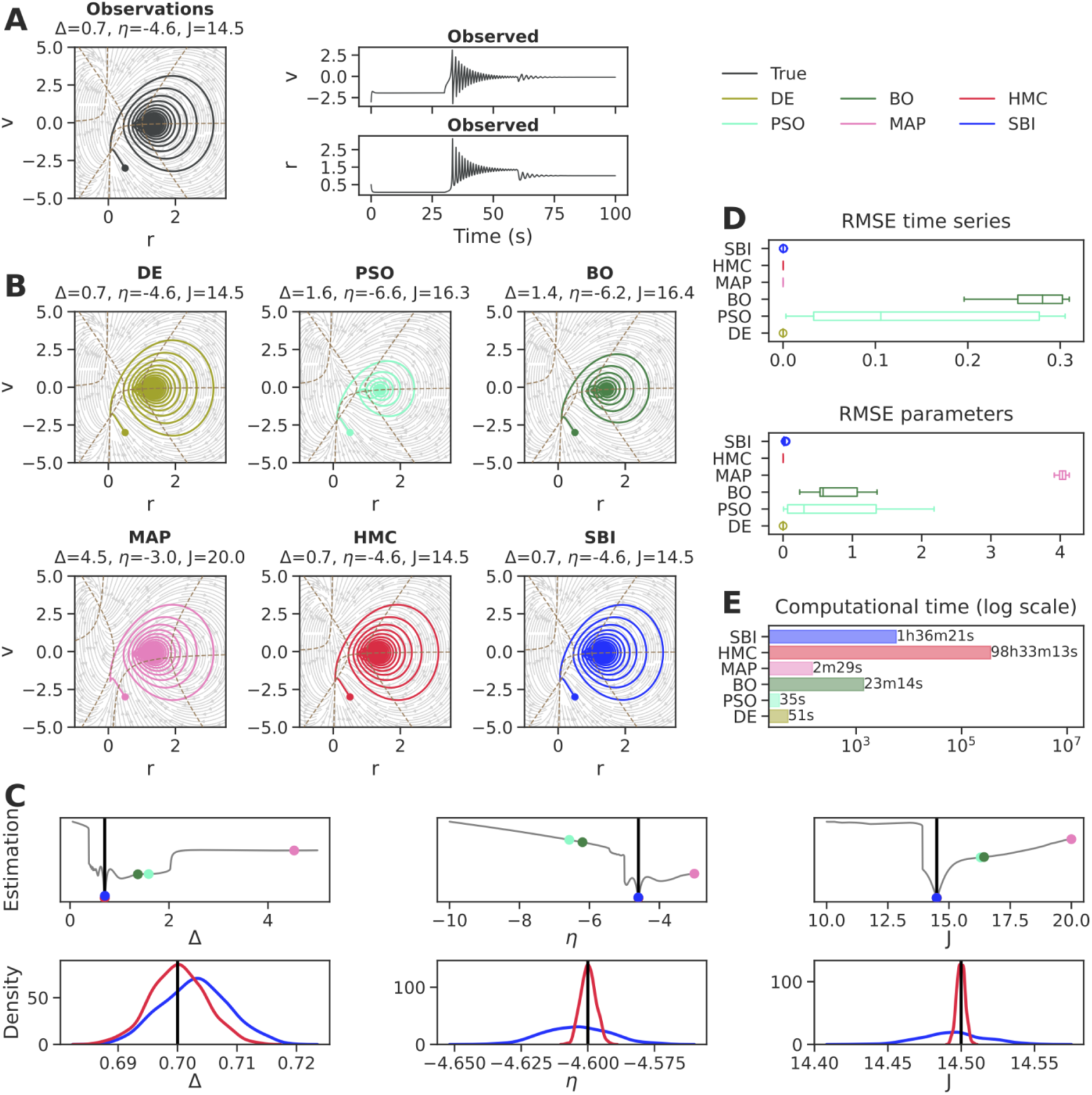
Inference on deterministic data. (**A**) The trajectories in phase-plane exhibiting bistable dynamics, and the corresponding time-series of firing rate (*r*) and membrane potential (*v*), used as the observations for inference (ground truth: Δ = 0.7, *η* = *−*4.6, *J* = 14.5). (**B**) The estimated trajectories in phase-plane, along with the corresponding parameters displayed in the top panels. All the algorithms capture the bistable dynamics in a qualitative manner. (**C**) For parameters Δ, *η*, and *J*, the point estimations are displayed in the top panels along with the profile likelihood (in grey), while the full posterior distributions are shown in the bottom panels. HMC leads to more precise parameter estimates with lower uncertainty compared to SBI. (**D**) Accuracy in estimation based on the sum over RMSE values. The bootstrap uncertainty is calculated for time-series (top panel) and parameters (bottom panel) through multiple runs. (**E**) Computational cost for each inference algorithm. Overall, DE excels in both speed and accuracy, but it offers only a point estimate. HMC is exact and provides informative posterior estimates, but it is very computationally prohibitive. Rather, SBI effectively provides accurate estimates along with associated uncertainties at a reasonable computational cost. DE, Differential Evolution; PSO, Particle Swarm Optimization; BO: Bayesian Optimization; MAP, Maximum a Posteriori; HMC: Hamiltonian Monte Carlo; SBI: Simulation-Based Inference; RMSE: Root Mean Square Error.

When it comes to matching the observed time-series, the optimization algorithms such as PSO and BO deviate significantly from true values, while other approaches equally retrieve an almost perfect fit to the observed time-series (see Figure 1**D**, top panel). Nevertheless, MAP largely fails to accurately estimate model parameters due to overfitting (Figure 1**D**, bottom panel). This type of overfitting emphasizes the importance of quantifying uncertainty to verify the reliability of inference, going beyond a point estimation. Note that for HMC and SBI, we report the results using posterior predictive check i.e., re-generating data using random parameters drawn from the estimated posterior and then comparing simulations with the observed data.

In terms of computational cost for inference, DE has a clear advantage with its rapid performance, typically taking less than a minute to complete (Figure 1**E**). Despite a high precision, the computational cost of HMC in this example is prohibitively expensive, taking almost a hundred hours to complete the inference process. On the other hand, SBI was terminated in nearly one and a half hours (including 100k random simulations, training, and sampling), making it approximately 60 orders of magnitude faster than HMC. Additionally, due to the amortized approach adopted by SBI, each sampling process takes less than one minute to estimate the joint posterior distributions from new data.

In summary, DE demonstrates superior speed and accuracy, but it only provides a point estimate. Among Bayesian methods, HMC is exact and provides certain estimates, but it is computationally prohibitive. On the other hand, SBI effectively provides accurate estimates along with associated uncertainties at a reasonable computational cost (see Table 1, [Movie 1](https://github.com/ins-amu/Inference_MFM/blob/main/Videos/Movie1_MPR_ODE_HMC_RV.mp4), and [Movie 2](https://github.com/ins-amu/Inference_MFM/blob/main/Videos/Movie2_MPR_ODE_SBI_RV.mp4)).

**Table 1.**
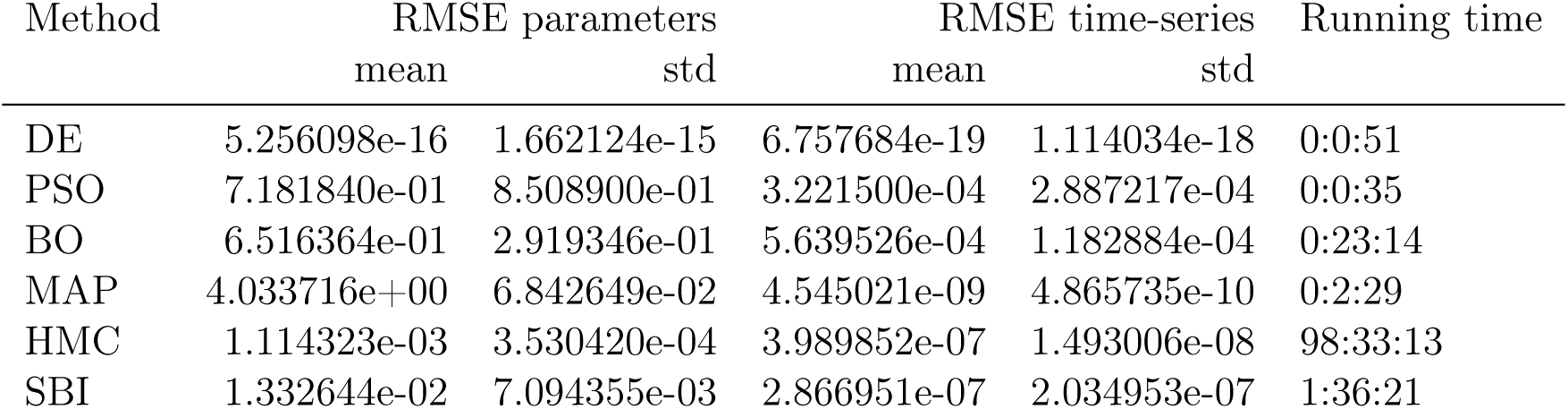
Benchmark on deterministic data.

| Method | RMSE parameters |  | RMSE time-series |  | Running time |
| --- | --- | --- | --- | --- | --- |
|  | mean | std | mean | std |  |
| DE | 5.256098e-16 | 1.662124e-15 | 6.757684e-19 | 1.114034e-18 | 0:0:51 |
| PSO | 7.181840e-01 | 8.508900e-01 | 3.221500e-04 | 2.887217e-04 | 0:0:35 |
| BO | 6.516364e-01 | 2.919346e-01 | 5.639526e-04 | 1.182884e-04 | 0:23:14 |
| MAP | 4.033716e+00 | 6.842649e-02 | 4.545021e-09 | 4.865735e-10 | 0:2:29 |
| HMC | 1.114323e-03 | 3.530420e-04 | 3.989852e-07 | 1.493006e-08 | 98:33:13 |
| SBI | 1.332644e-02 | 7.094355e-03 | 2.866951e-07 | 2.034953e-07 | 1:36:21 |

### 3.2. Inference on stochastic data

The inherent randomness in stochastic systems introduces uncertainty into the behavior of the system under study, which makes it challenging to accurately identify its underlying dynamics. In particular, the presence of dynamical noise significantly increases the complexity of the inference process and necessitates the use of robust probabilistic methodologies that can effectively account for and handle the stochastic nature of the data.

Here, we report the results of inference on synthetic data with zero-centered Gaussian dynamical noise, where the intensity is *σ* = 0.1 (see Figure 2). The observed noisy time-series and trajectories in phase-plane, exhibiting a bistable behavior between a stable node and a stable focus, are illustrated in Figure 2**A**. From the results demonstrated in Figure 2**B**, we can see that all the algorithms qualitatively replicate the bistability behavior in the phase-plane. Yet, when assessing how well they recover the true parameters, only DE yields an accurate point estimate among the optimization algorithms (Figure 2**C**, top panels).

**Figure 2.**
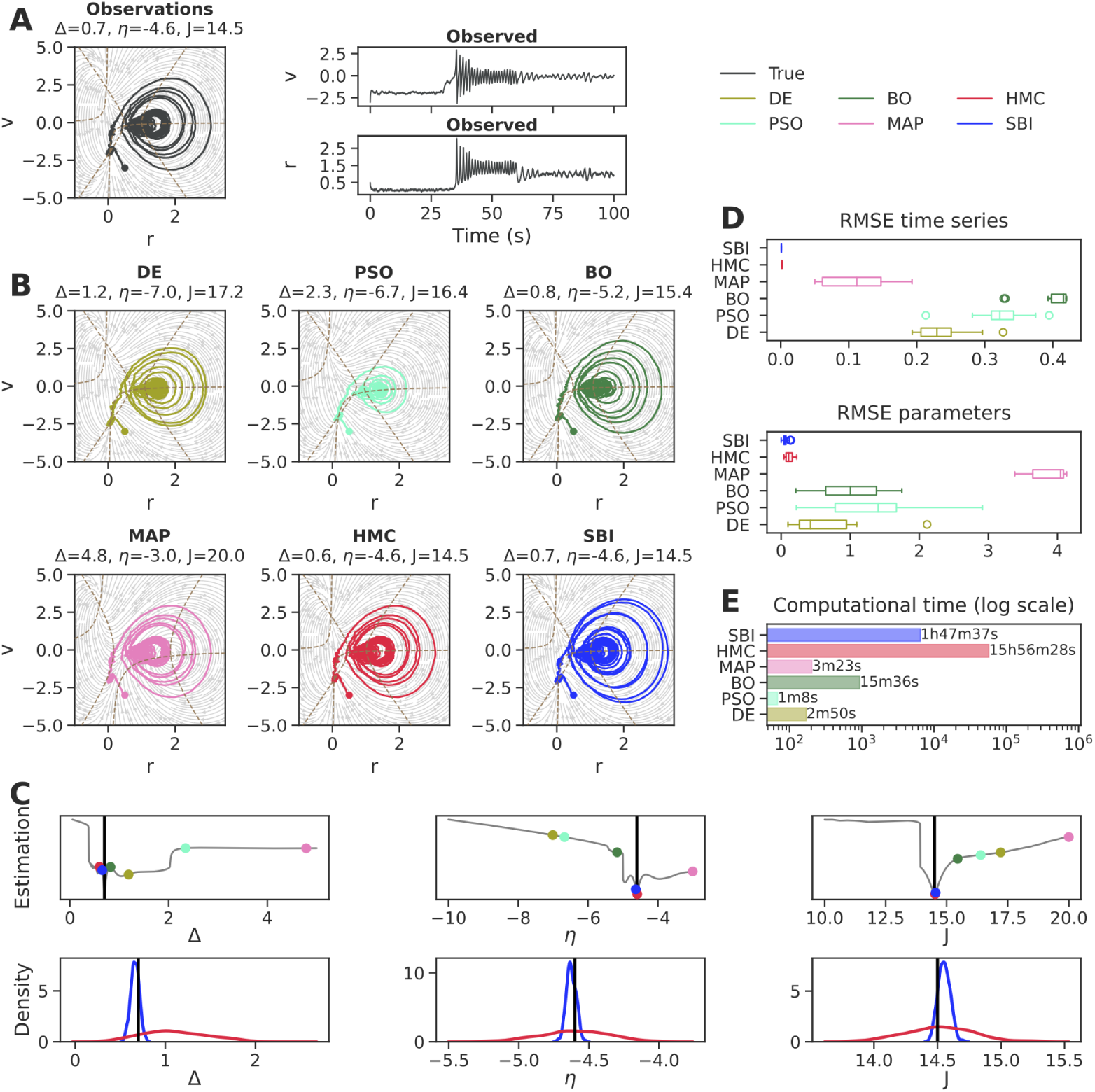
Inference on stochastic data. (**A**) The generated bistable dynamics in the phase-plane and the corresponding time-series of firing rate (*r*) and membrane potential (*v*) used as the observations for inference (ground truth: Δ = 0.7, *η* = *−*4.6, *J* = 14.5). (**B**) The estimated trajectories in the phase-planes, along with the corresponding parameters displayed in the top panels. All the algorithms qualitatively capture the bistable behavior. (**C**) The point estimation along with the profile likelihood (top panels) and the full posterior (bottom panels) for parameters Δ, *η*, and *J*. SBI leads to more precise parameter estimates with lower uncertainty compared to HMC. (**D**) Accuracy in estimation based on the sum over RMSE values for the time-series (top panel) and parameters (bottom panel). The bootstrap uncertainty is calculated through multiple runs. (**E**) Computational cost for each inference algorithm. When evaluating overall accuracy, uncertainty quantification, and computational cost, then SBI outperforms all other algorithms.

In terms of uncertainty quantification, SBI generates posteriors that are tightly centered on the ground-truth parameters, in comparison to the HMC sampling (see Figure 2**C**, bottom panels, and Figure S5). A detailed comparison of convergence diagnostics indicates that both SBI and HMC methods provide ideal Bayesian estimation (see Figure S6). Regarding the proximity to the observed time-series and true parameters, both HMC and SBI offer a closer match compared to the other algorithms (Figure 2**D**). This highlights the challenges inherent in inferring stochastic systems via optimization algorithms, as the error metrics such as RMSE used to define the objective function may not reliably gauge accuracy.

In terms of computational efficiency for inference, the running time of all algorithms increases when the noise is present, except for HMC (Figure 2**E**). Nevertheless, when considering the entire process (including random simulations, training, and sampling), SBI remains approximately 8 orders of magnitude faster than HMC. Interestingly, SBI is also able to accurately and efficiently estimate the dynamical noise in the system (see Figure S7).

In summary, these results demonstrate that when considering overall accuracy in the presence of noise and bistability, uncertainty quantification, and computational cost, the SBI outperforms all other algorithms, including HMC (see Table 2, [Movie 3](https://github.com/ins-amu/Inference_MFM/blob/main/Videos/Movie3_MPR_SDE_HMC_RV.mp4), and [Movie 4](https://github.com/ins-amu/Inference_MFM/blob/main/Videos/Movie4_MPR_SDE_SBI_RV.mp4)).

**Table 2.** Benchmark on stochastic data.

| Method | RMSE parameters |  | RMSE time-series |  | Running time |
| --- | --- | --- | --- | --- | --- |
|  | mean | std | mean | std |  |
| DE | 0.672351 | 0.617302 | 0.000533 | 9.850079e-05 | 0:2:50 |
| PSO | 1.362540 | 0.852908 | 0.000717 | 1.116449e-04 | 0:1:8 |
| BO | 1.021078 | 0.519784 | 0.000888 | 8.232933e-05 | 0:15:36 |
| MAP | 3.883882 | 0.302616 | 0.000250 | 1.228436e-04 | 0:3:23 |
| HMC | 0.121416 | 0.081035 | 0.000004 | 1.427944e-08 | 15:56:28 |
| SBI | 0.052948 | 0.024310 | 0.000002 | 2.178518e-07 | 1:47:37 |

Next, we investigate how varying the intensity of dynamical noise impacts the inference on stochastic data. Our results indicate that the overall quality of fit to the noisy data (Figure 3**A**) and the accuracy of recovered parameters (Figure 3**B**) remain stable up to σ ≤ 10^−1^ for all algorithms. However, both metrics exhibit a significant increase beyond this threshold, particularly for optimization methods.

**Figure 3.**
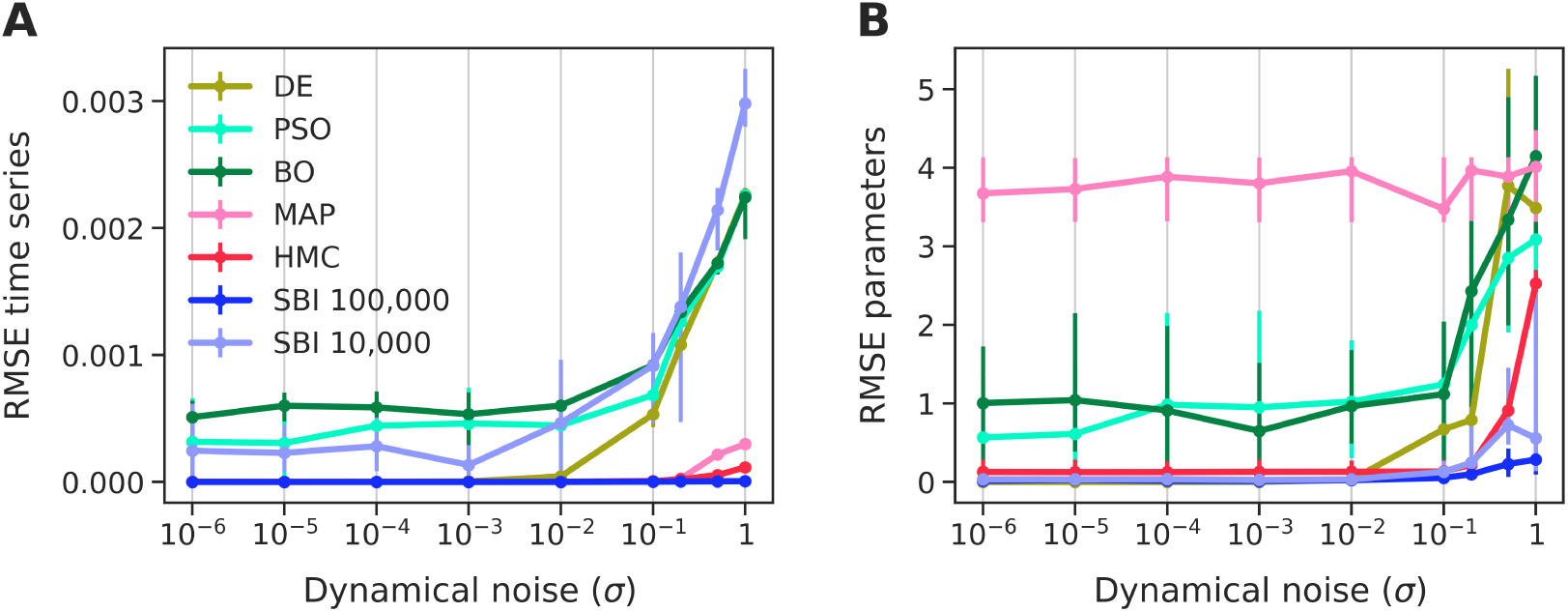
The performance of inference algorithms with increasing the intensity of dynamical noise using the sum over: (**A**) RMSE of the time-series, and (**B**) RMSE of the model parameters. Overall, with a sufficient number of simulations for training, the SBI emerges as the most accurate and robust method in our algorithmic benchmark.

The MAP estimation is robust with regard to the amount of dynamical noise, but it tends to overfit. This is because the MAP estimation consistently provides the least accurate inference on parameters, even though its fit to the time-series data is almost perfect. In contrast, Bayesian inference methods such as HMC and SBI are overall more accurate and significantly more robust compared to optimization methods. Both HMC and SBI exhibit significantly lower error than others, even at high noise levels (e.g., *σ* = 1).

Interestingly, SBI appears to be more resilient to high levels of noise compared to HMC. Given a sufficient number of simulations for training (e.g., 100k), SBI demonstrates the most accurate and robust fit to the data, consistently staying close to a perfect fit across various noise values. See Figure S8 for a systematic investigation on the impact of the number of simulations on the performance of SBI. Overall, these results indicate that SBI is the most accurate and robust algorithm in our benchmark.

### 3.3. Inference on missing data in the state-space

Here, we explore the performance of inference methods in situations where data is available for only one of the variables in the state-space modeling. We consider the mean membrane potential *v* as the observed data, which is simulated with dynamic noise of intensity *σ* = 0.1, while the firing rate *r* remains latent (Figure 4**A**). The noise intensity is fixed for inference process.

**Figure 4.**
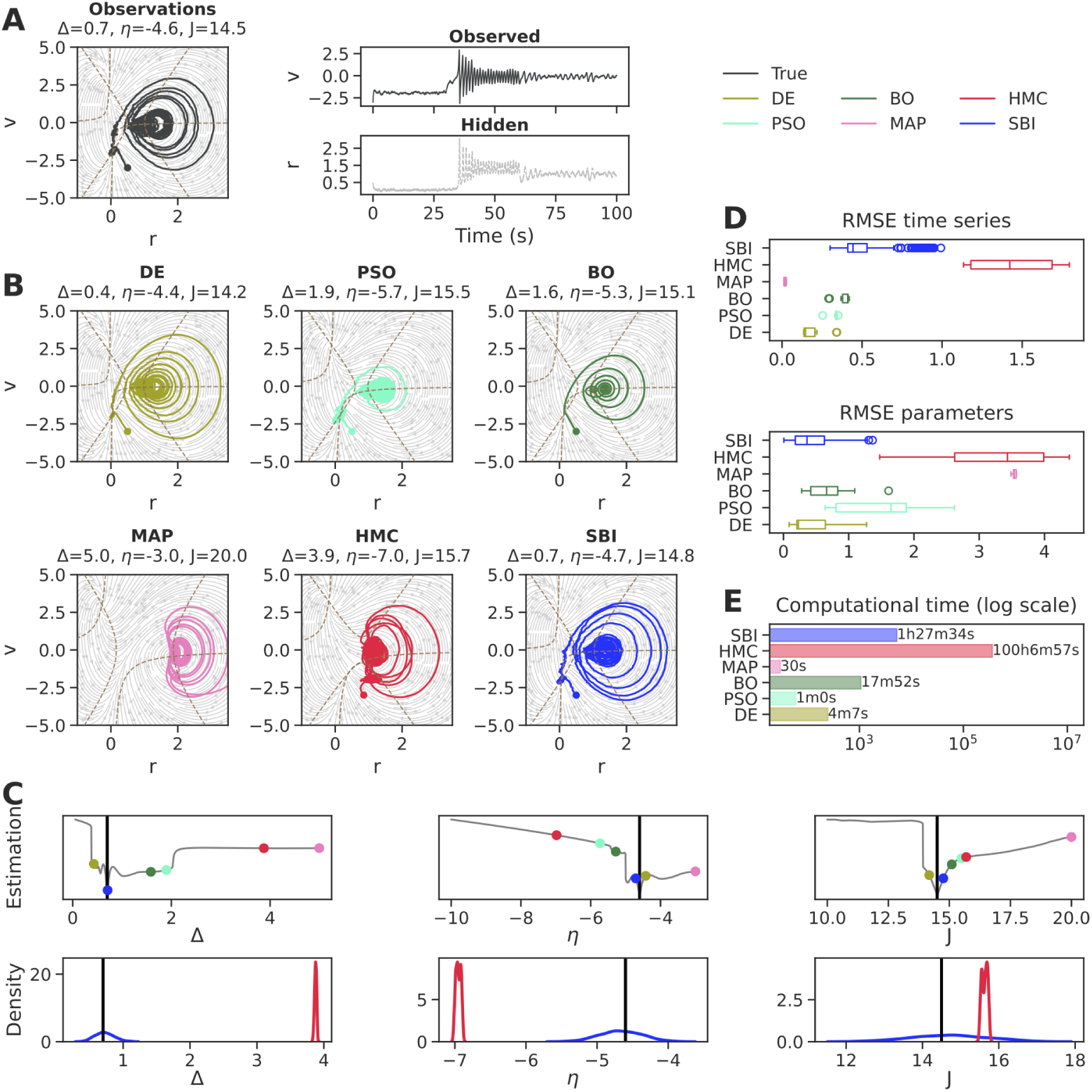
Inference on noisy observation with missing data. The generated bistable dynamics in the phase-plane and the corresponding time-series of observed membrane potential (*v*) and hidden firing rate (*r*), with ground truth: Δ = 0.7, *η* = *−*4.6, *J* = 14.5. (**B**) The estimated trajectories in the phase-planes, along with the corresponding parameters displayed in the top panels. Only DE and SBI provide a close agreement with the observed bistable trajectories. (**C**) The point estimation along with the profile likelihood (top panels), and the full posterior (bottom panels) for parameters Δ, *η*, and *J*. (**D**) The accuracy of estimation is evaluated by summing the RMSE values for both the true time-series (top panel) and parameters (bottom panel), while considering the bootstrap uncertainty through multiple iterations. (**E**) The computational cost for each inference algorithm. Overall, DE and SBI outperform other algorithms in inferring the bistable dynamics when dealing with missing data.

From Figure 4**B**, we observe that only DE and SBI were capable of retrieving the true dynamics in the phase-plane when *r* was missing. As shown in Figure 4**C** (top panel), among optimization algorithms, only DE correctly estimates the true parameters. Notably, HMC fails considerably in this problem by proposing over-confident posterior distributions that are far from the ground truth (Figure 4**C**, bottom panel). In contrast, the estimated posteriors using SBI are centered on the ground-truth parameters, but they exhibit a more diffuse uncertainty compared to the previous results (see Figure S9). Note that HMC fails to accurately reconstruct the latent variable *r* from observed *v* (see Figure S10), even though its error on the fitted trajectories is comparable to that of SBI (Figure 4**D**, top panel). Nevertheless, the unreliability in the results produced by HMC becomes evident when observing the RMSE values for model parameters (Figure 4**D**, bottom panel).

In terms of computational cost for inference (Figure 1**E**), optimization by DE still has a clear advantage with its rapid performance. Interestingly, SBI remains efficient, approximately 68 orders of magnitude faster than HMC (see Table 3, [Movie 5](https://github.com/ins-amu/Inference_MFM/blob/main/Videos/Movie5_MPR_SDE_HMC_V.mp4), and [Movie 6](https://github.com/ins-amu/Inference_MFM/blob/main/Videos/Movie6_MPR_SDE_SBI_V.mp4)). Overall, SBI outperforms HMC in the recovery of the bistable dynamics, including the hidden firing rate. However, this approach can lead to an overestimation in the associated uncertainty, when compared to the full observed state-space dynamics (see Figure 2 versus Figure 4).

**Table 3.**
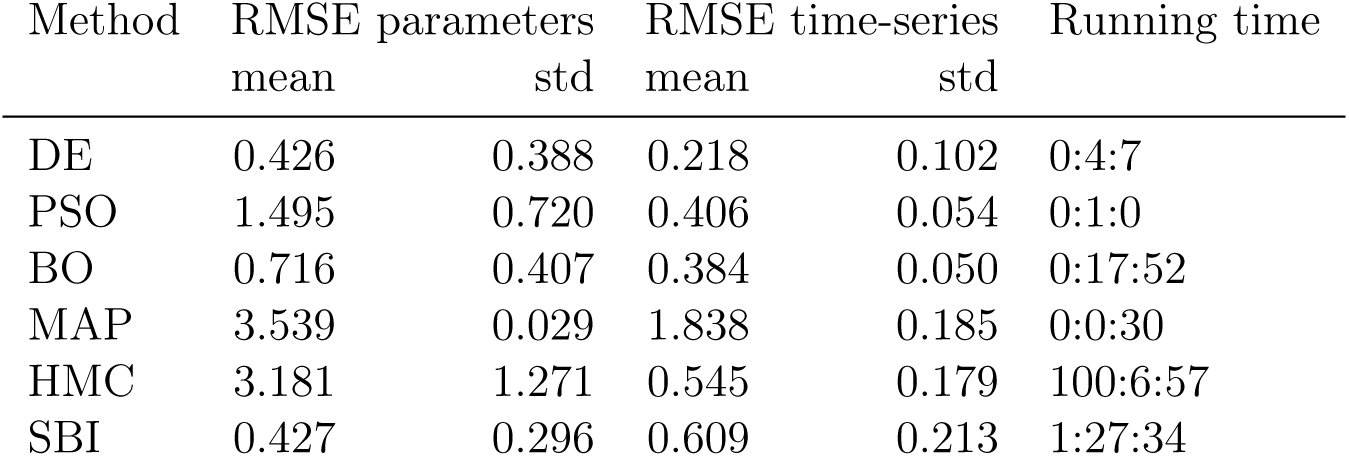
Benchmark on missing data.

### 3.4. Phase-space reconstruction

As demonstrated in the previous section, the inference process posed a challenge when only the membrane potential *v* was observed and the firing rate *r* was missing. To improve the inference in such cases, we approximately reconstruct the hidden *r* from the observed counterpart *v*, using the time-delay embedding technique (Figure 5).

**Figure 5.**
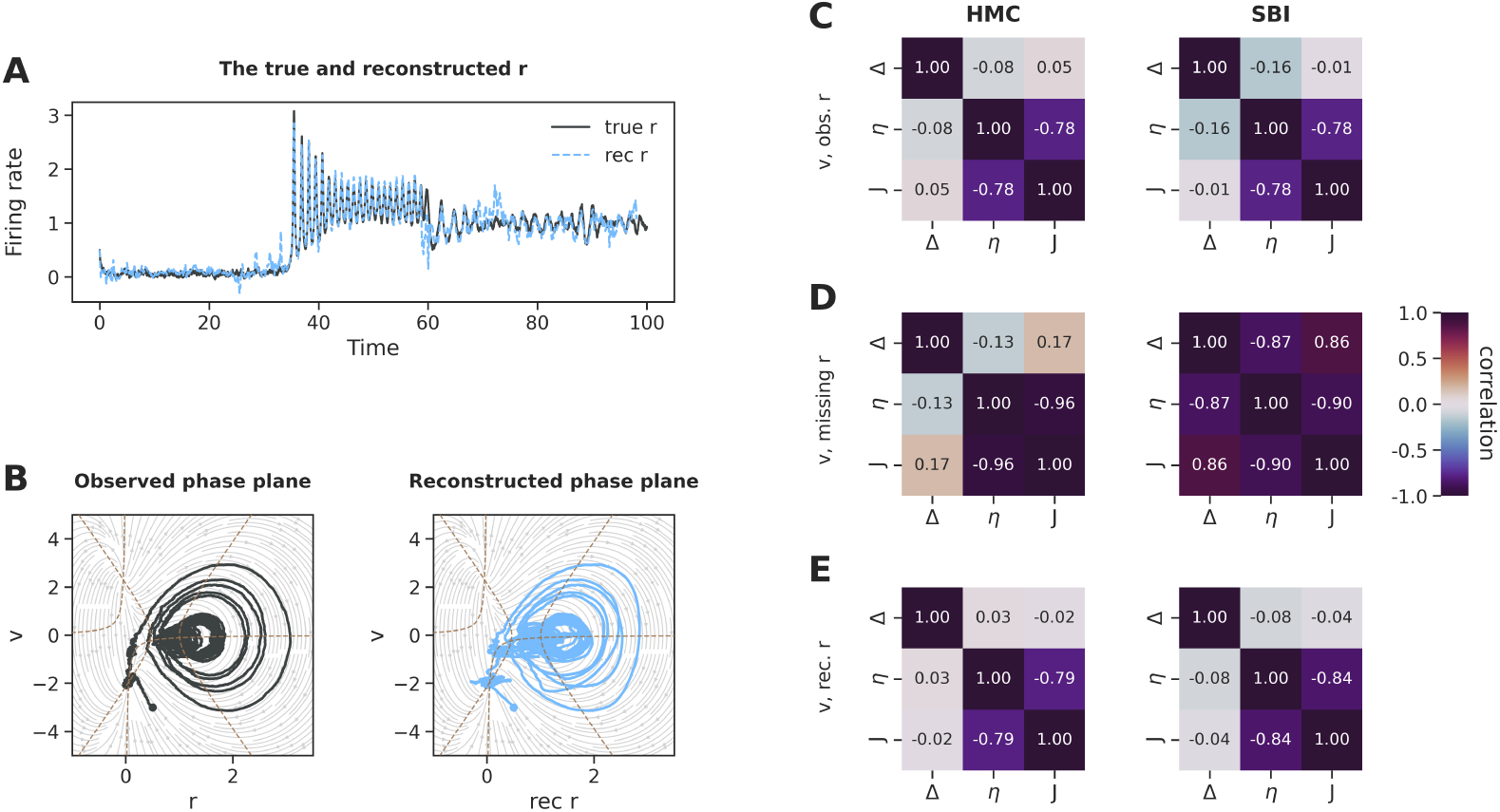
Reconstruction of firing rate *r* from mean membrane potential *v*, and comparison of inter-dependency between parameters. (**A**) Original and reconstructed mean firing rate. (**B**) Observed (left) and reconstructed (right) system dynamics in phase-plane. Correlations between estimated joint posterior distributions, using HMC (left) and SBI (right), when: (**C**) Both *r* and *v* are observed, (**D**) Only *v* is observed and *r* is hidden, (**E**) Reconstructed *r* from the observed *v* using time-delay embedding.

As it can be seen from Figure 5**A**, the reconstructed firing rates closely follow the original time-series (RMSE=0.151). This leads to accurately capturing the bistable switching behavior in the phase-plane, as shown in Figure 5**B**. We subsequently explored how the access to the trajectories of state variables influences the statistical relationship between parameters. When both the firing rate (*r*) and membrane potential (*v*) are observed, both HMC and SBI algorithms reveal a strong correlation between the average excitability *η*, and the synaptic weight *J* (*ρ_η,J_ ≈ −*0.78), while the other parameters exhibit no codependency (see Figure 5**C**).

When the firing rate is latent and only the membrane potential is observed, the correlation between parameters is more pronounced, as estimated by HMC, while SBI tends to overestimate this co-dependency as fully degenerate (see Figure 5**D**). This highlights the challenges in the inference or parameter estimation process when lacking access to complete knowledge of the system dynamics. This issue can be improved by using time-delay embedding to reconstruct the full phase-space dynamics and inform the inference process, as shown by the reduced correlation between parameters in Figure 5**E**.

### 3.5. Inter-dependency between generative parameters

In the previous sections, we performed inference against the system dynamics derived from the MF model given by Eq. (1). We now aim to investigate how the inference of the posterior distribution and the relationships between parameters remains consistent by traversing across scales. This can be achieved by conducting inference using observed data generated by QIF neurons (at the microscopic level) versus the data generated by MF model (at the macroscopic level).

First, we compared the simulated membrane potential *v* and firing rate *r* using a MF model given by Eq. (1) to the averaged activities of a network of 10^4^ all-to-all connected QIF neurons (see raster plot shown in Figure 6(**A**)). Figure 6(**B**) demonstrates that the MF model can accurately generate the (smoothed) transient dynamics that emerge from an ensemble of spiking neurons. However, the first spike emitted by the MF model after stimulation may exhibit a short lag compared to the averaged QIF neurons. Then, we used the data generated by MF and QIF models as the observation, and conducted inference using the MF model through Bayesian estimation algorithms, such as HMC and SBI. Specifically, we compared the correlations between the estimated joint posteriors of model parameters in each case, as shown in Figure 6(**C**).

**Figure 6.**
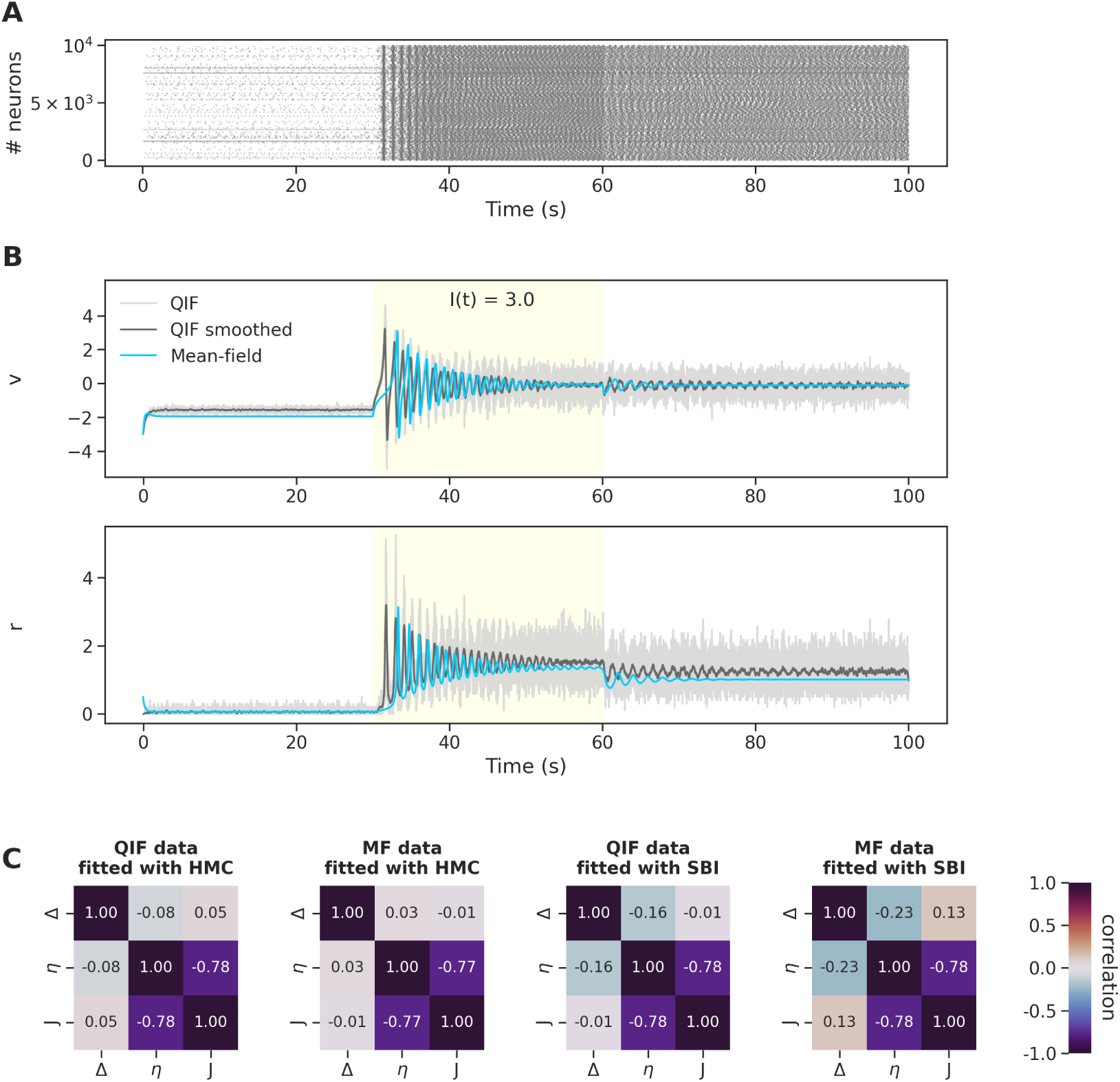
Comparing the transient dynamics of a MF model with the emergent dynamics of a network of QIF neurons, and exploring the interdependency between parameters. (**A**) Raster plot of QIF neurons. (**B**) The membrane potential *v* (top panel) and firing rate *r* (bottom panel) generated by MF model (in cyan) versus averaged activities of QIF neurons, as the raw simulations (in light gray) and smoothed (dark grey). At time *t* = 30 *sec*, a current *I*_0_ = 3 *mv* is applied to all neurons, and set to zero again at *t* = 60 *sec*. (**C**) The Pearson correlation coefficients between parameters in the MF model, estimated using Bayesian algorithms (HMC and SBI), against data generated by different models (MF and QIF). The strong negative linear correlation between excitability and synaptic weight (*ρ_η,J_ ≈ −*0.78) persists across algorithms and datasets.

We observed consistent correlations between parameters across the two types of datasets (macroscopic MF and microscopic QIF) when fitted using the MF model. Our results indicate that the strong negative linear correlation between excitability and synaptic weight (*ρ_η,J_ ≈ −*0.78) persists when using HMC and SBI against both datasets. Considering the agreement between HMC and SBI across two datasets, we can conclude that the consistent high correlation between parameters *η* and *J* is intrinsic and not induced by the inference process or model assumptions.

### 3.6. SBI on the stimulus current

Here, we challenge SBI approach in inferring system dynamics while also accounting for an unknown input current, which plays a crucial role in emerging the bistable behavior. Given only the position or waveform of input currents, we estimate the intensity or angular velocity across various ground-truth values: *I*_0_ in a step current as *I*(*t*) = *I*_0_ *·* **1**_stim_(*t*) and *ω*_0_ in a sinusoidal current as *I*(*t*) = sin(*ω*_0_*t*) *·* **1**_stim_(*t*), as shown in Figure 7(**A**), and (**B**), respectively. Our results demonstrate that in both cases, the SBI approach accurately recovers the unknown parameters in the input currents. The posterior credibility intervals, visualized as error bars, indicate certain estimates that are close to a perfect fit (*y* = *x*, in red) across all different values. See Figure S11 for the observed and predicted time-series. This validates the capability of SBI in accurately estimating the system dynamics, even when the characteristics of the input current are unknown.

**Figure 7.**
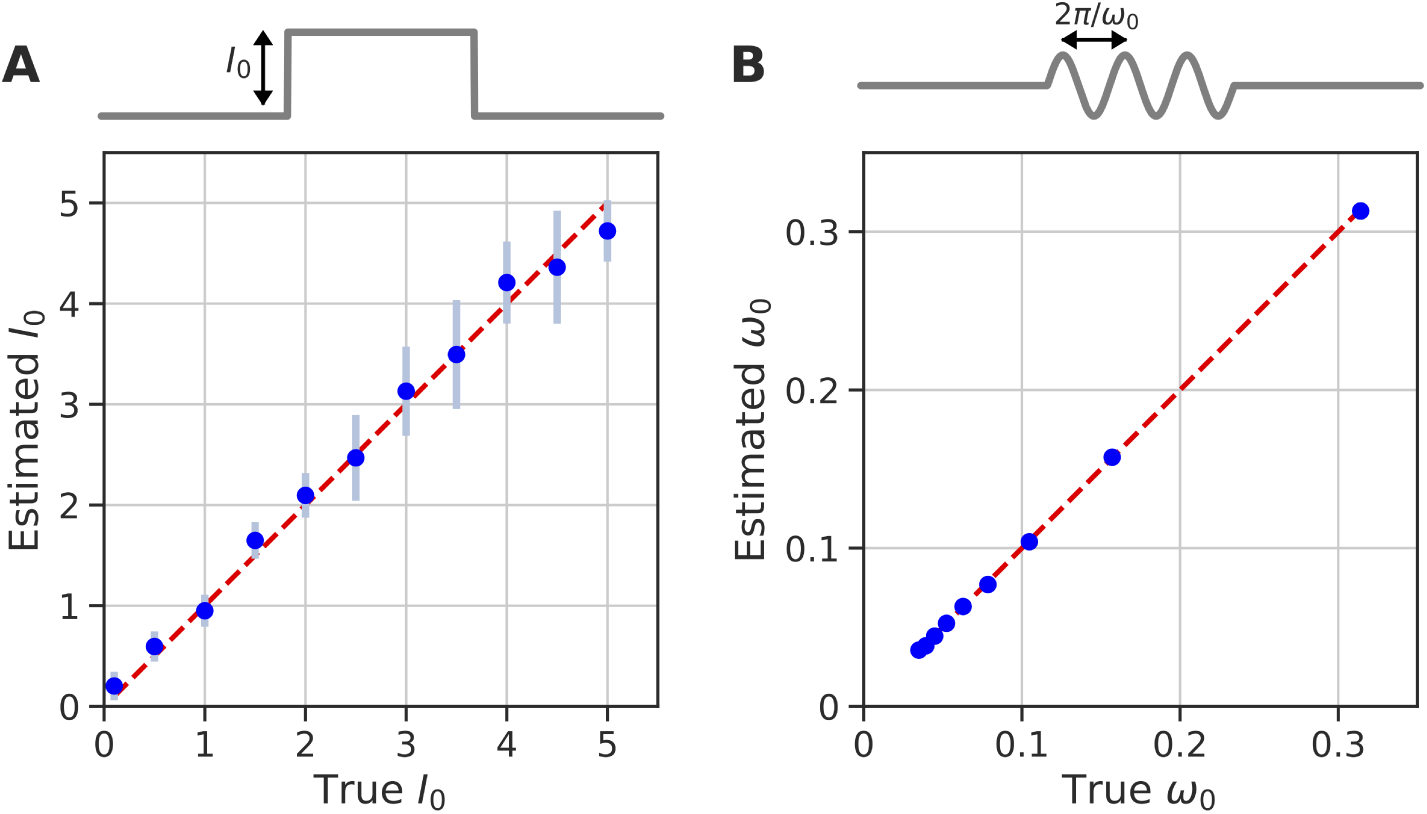
SBI on the stimulus current. (**A**) Step current as *I*(*t*) = *I*_0_ *·* **1**_stim_(*t*). (**B**) Sinusoidal current as *I*(*t*) = sin(*ω*_0_*t*) *·* **1**_stim_(*t*). Dashed red line represents a perfect fit.

### 3.7. SBI on stability of system dynamics

Here, we show that SBI can be used to investigate the stability of system dynamics from low-dimensional summary statics of observed time-series. Figure 8 shows a phase diagram of the system as a function of the mean *η* and synaptic weight *J*, both normalized by the width of the input distribution Δ. Using linear stability analysis, there are three qualitatively distinct regions of the phase diagram: (i) A single stable node corresponding to a low-activity state (shown in blue), (ii) A single stable focus generally corresponding to a high-activity state (shown in red), and (iii) A region of bistability between low and high firing rate (shown in cyan).

**Figure 8.**
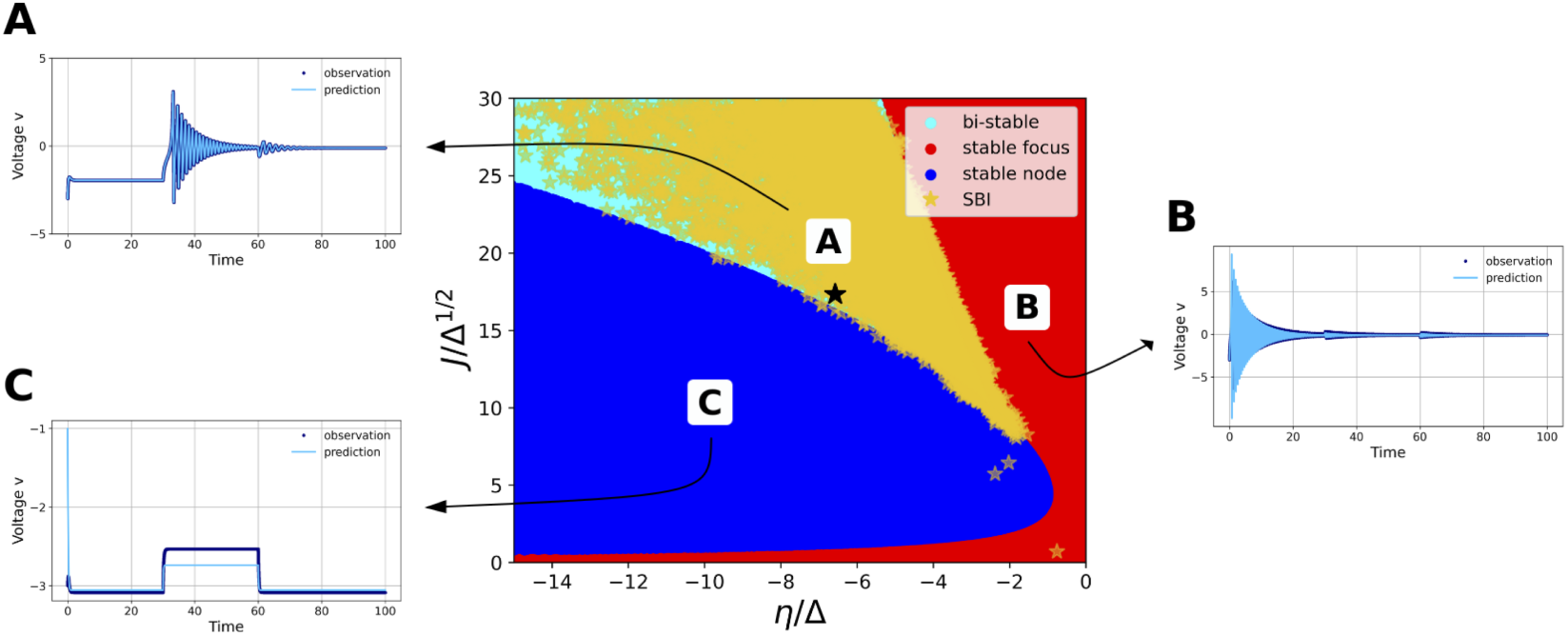
SBI over the stability of system dynamics in phase diagram. In the wedge-shaped region (shaded in cyan), bistability exists between a high and a low activity state. On the right side, a stable focus is indicated (shaded in red), while on the left side, a stable node is depicted (shown in blue). By training a deep neural density estimator on data features, the posterior samples generated using SBI accurately capture the bistable dynamics (shown in yellow), aligning closely with results from linear stability analysis. The black asterisk denotes the observation point used for inference. The insets display observed (dark blue) and predicted (light blue) time-series in different regimes: (**A**) Bistability, (**B**) Stable focus, and (**C**) Stable node.

Interestingly, a similar basin of bi-stability in phase diagram can be readily reproduced using deep neural density estimators (such as MAF model) in SBI approach. By training on the low-dimensional data features extracted from the time-series (such as the presence or absence of damped oscillations before, during, and after stimulation), the generated posterior samples display a very close agreement with the results obtained from linear stability analysis. This demonstrate the capability of SBI in accurately estimating system dynamics from summary statics, including the presence of bi-stability in the phase diagram.

### 3.8. Neural ODEs on system dynamics

In this section, our aim is to infer the collective dynamics of QIF neurons without making any assumptions about the underlying generating dynamics. To achieve this, we used Neural ODEs as a powerful tool for modeling continuous-time dynamics without assuming any prior knowledge of the underlying equations governing the system. Unlike traditional discrete-time models, Neural ODEs parameterize the continuous-depth formulation, allowing for seamless interpolation between observed data points.

We first validate Neural ODEs using the data garnered by MF model described by Eq. (1). We generated 3 different datasets using the MF model given by Eq. (1), with a set of parameters corresponding to a bistable regime: {Δ = 1, *J* = 15, *η* = *−*5}. For each datasets, we sampled the phase-space by varying initial conditions according to a regular grid so that *r*_0_ ∈ [0.1, 3] and *v*_0_ ∈ [*−*2, 2]. We then solved the system for 1000 time points (with an integration time step of *dt* = 0.01 *sec*). To speed up training, each trajectory was downsampled by a factor of 10 and divided into 10 segments, each consisting of 10 time points. The training and test data were randomly split, with 75% of the data used for training and 25% used for testing.

The first training set, comprising of *N* = 100 deterministic trajectories, was generated (see Figure 9**A**). The results indicate that the Neural ODE almost perfectly reconstructed the phase-space (see Figure 9**B**). In the second training set, which had the same size, a dynamical noise with a standard deviation of *σ* = 0.1 was added during the integration process (Figure 9**C**). In this case, the estimated nullclines suffered from overfitting although without affecting the overall reconstructed dynamics, as the predicted trajectories were still very similar to the original data (as shown Figure 9**D**). Increasing the size of the training dataset (see Figure 9**E**) significantly reduces overfitting. As a result, the reconstructed nullclines show a very close agreement with those obtained from the deterministic data (Figure 9**F**). To illustrate overfitting, Figure S12 shows the loss functions for the different scenarios, as well as snapshots of phase-space reconstruction during training. Overall, the Neural ODE successfully reconstructs the underlying deterministic system, even in the presence of noise (see [Movie 7](https://github.com/ins-amu/Inference_MFM/blob/main/Videos/Movie7_MPR_PhasePlane_NeuralODEs.mp4)).

**Figure 9.**
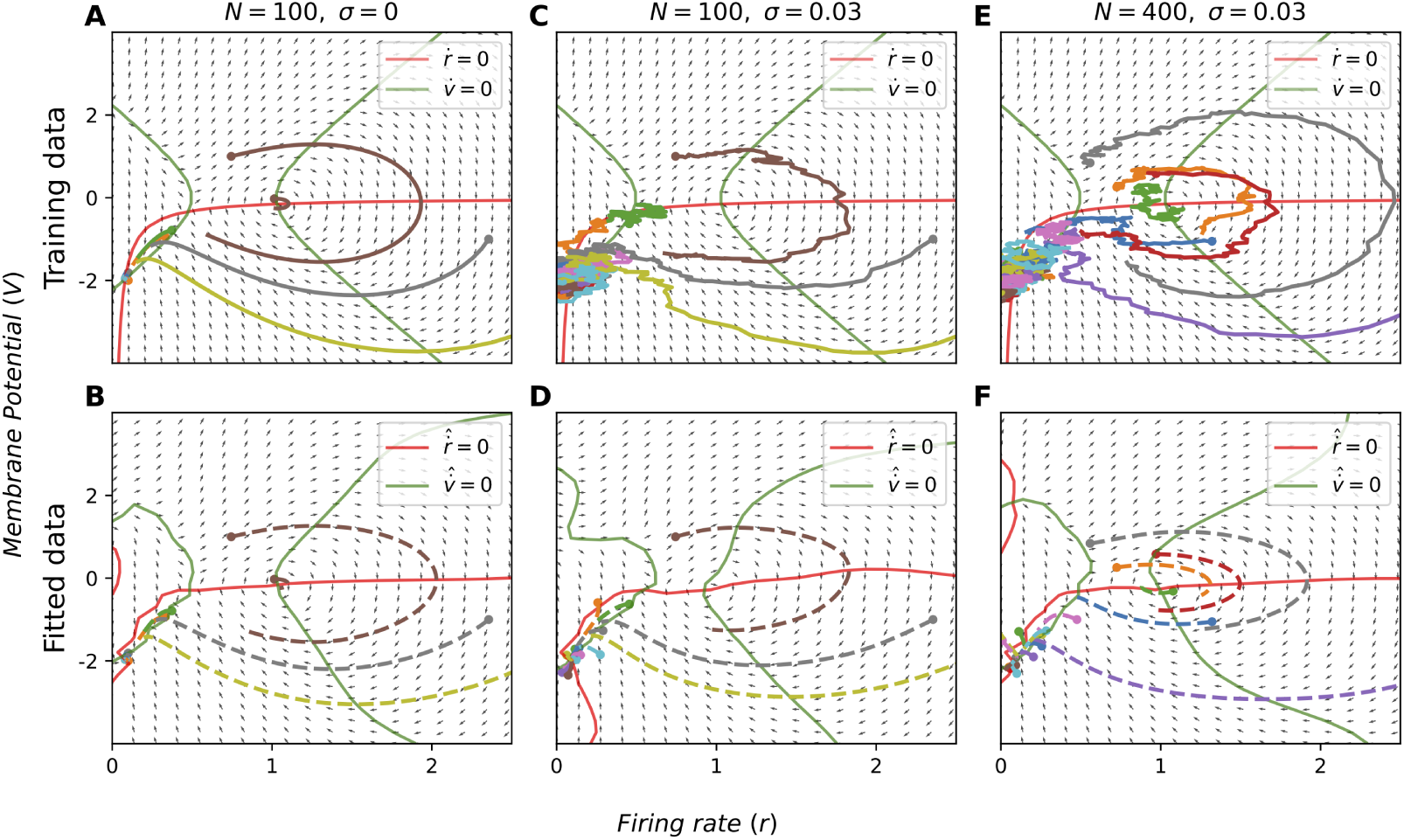
Example of training data generated from the MF model set in a bistable regime (top row). The trajectories are displayed after the 10-fold split, without the downsampling step for better visualization. Corresponding fit obtained with the Neural ODE for the same initial conditions and estimated phase-plane (bottom row) after 45000 training iterations. Either (**A**, **B**) using a training set of 100 deterministic trajectories, (**C**, **D**) using a training set with dynamical noise, (**E**, **F**) or a larger dataset. The red and green curves represent the nullclines of *r* and *v*, respectively. The dots represent the initial values for each trace, while the dashed lines correspond to the Neural ODE trajectories. The neural ODE is prone to overfitting when noise is introduced in the training data, although it still preserves the overall dynamics. A larger dataset helps in recovering smoother nullclines.

We then trained Neural ODEs using data generated by 10^4^ QIF neurons with a uniform stimulus (see Figure 10). The data was partitioned using the first 400 points for training and predicting the remaining 1600 points. The results indicate that using derivatives dynamics, we can achieve a reliable understanding and prediction of the complex behavior of a network of spiking neurons, as illustrated in Figure 10. The emergent dynamics vary based on different parameter settings, resulting the stable node, stable focus, and bistable regime. See Figure S13 for the loss function in the training and test sets.

**Figure 10.**
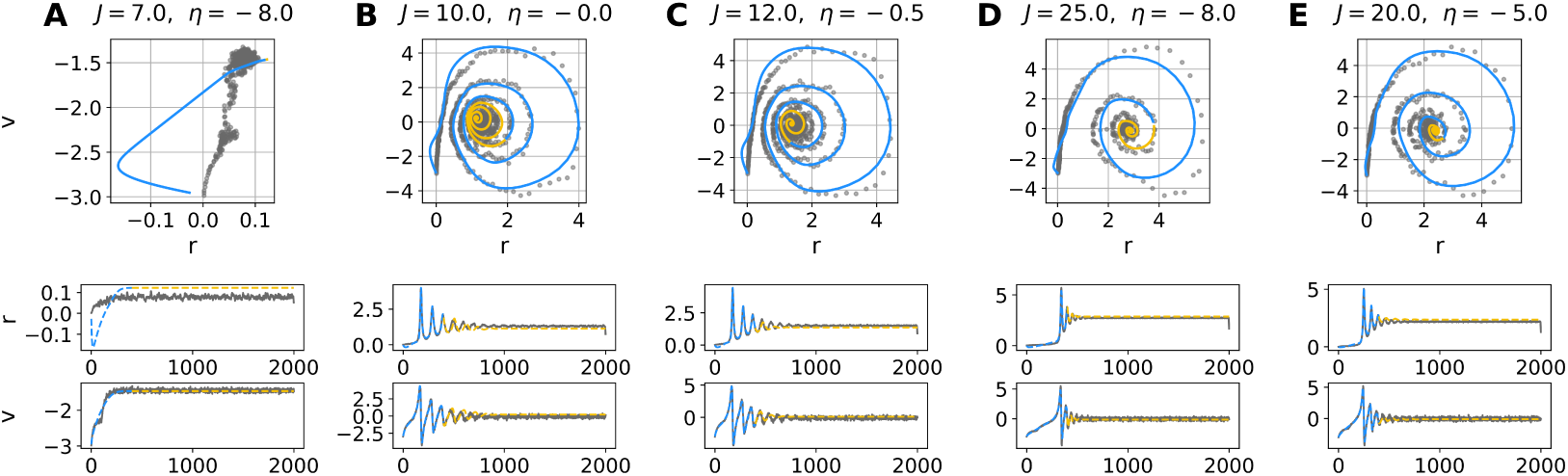
Reconstruction and extrapolation on QIF data with various parameters and dynamic regimes, using Neural ODEs. (**A**) Stable nodes, (**B**, **C**) Stable foci, (**D**, **E**) Bistability. The grey dots represent sparse observations, blue lines represent predictions (400 points), and orange lines represent extrapolations (1600 points). We can see that our Neural ODE has learned the system dynamics and provides accurate predictions based on data generated at the microscopic level.

## 4. Discussion

In the ever-evolving field of computational neuroscience, accurately estimating parameters that consistently govern the collective behavior of neural networks is a crucial endeavor, especially within the framework of recurrently coupled spiking neurons. In this study, we emphasize the use of mean-field (MF) theory to streamline the inference process for networks of spiking neurons. This choice is driven by the computational challenges in the calculation of the likelihood function—an essential ingredient for both frequentist and Bayesian inference methods—that becomes computationally prohibitive when attempting inference using these spiking networks in forward modeling.

The needs and objectives of researchers in the field of computational neuroscience are as diverse as the neural systems they aim to understand. Some studies may prioritize rapid point estimation through optimization, seeking quick outcomes to inform real-time decisions (Vattikonda et al., 2021; Penas et al., 2023). On the other hand, some studies find value in exploring the full distribution of parameters, which provides a nuanced understanding of parameter uncertainty for reliable decision making (Hashemi et al., 2020; Jha et al., 2022). This paper offers a comprehensive, yet non-exhaustive, benchmarking of the state-of-the-art inference methods applied to a MF model of spiking neurons (see Figure 1, Figure 2, and Figure 4). Our comparative analyses offer practical guidance, assisting researchers in selecting the most suitable method for their specific datasets and research inquiries. Note that the comparison of computational cost and accuracy in parameter estimation can heavily depend on the selection of hyperparameters, such as population size in DE, warmup phase in HMC, or the number of simulations and features in SBI. To make an unbiased assessment, we used optimal values for these hyperparameters in each algorithm, tailored to the dynamical model used in this study. Nevertheless, these optimal values may vary for different inverse problems, depending on parameter space and data dimensions, nonlinearity, sparsity, and the mapping function to measurements.

While the optimization method can construct a confidence interval for the estimation based on a threshold for accepting or rejecting the estimates, the results are highly dependent on the chosen threshold value (Beaumont et al., 2002; Cranmer et al., 2020). In contrast, Bayesian inference naturally provides uncertainty quantification by placing a distribution over parameters and treating them as random variable. Following Bayes’ rule, this distribution is updated with evidence from observed data to form the posterior distribution, which furnishes comprehensive information for inference and prediction.

Our results indicated that evolutionary algorithm for solving global optimization problems, such as DE, provide rapid and accurate point estimation of the true generative parameters when there is no dynamical noise present. Challenges arise when using optimization methods in the presence of noisy data, mainly due to the lack of an efficient form of the objective function. The selection of the objective function plays a crucial role in determining the estimation through optimization methods (Svensson et al., 2012; Hashemi et al., 2018). The error explanation with distance metrics such as RMSE is limited in accurately capturing the underlying data generation process (Baldy et al., 2023). This is because when generative parameters remain unchanged but when dynamic noise is introduced, the time-series can show large fluctuations, resulting in deviations from the observed data. In particular, the presence of noise can easily lead to unreliable estimations, increasing the risk of overfitting, where the model fits to the noise rather than capturing the true underlying relationship. Consequently, it becomes more conspicuous to conduct inference using distributions in the Bayesian framework, particularly when dealing with dynamical noise. Diagnosing of overfitting, as shown using MAP estimation (see Figure 2 and Figure 4), can be better understood through uncertainty quantification.

Furthermore, Bayesian inference reveals the relationships between parameters, capturing degeneracy in the data (Edelman and Gally, 2001; Hashemi et al., 2023). For example, when we assess the agreement between HMC and SBI on different datasets (as shown in Figure 5, Figure S4, Figure S5, and Figure S9), it can be concluded that the persistent strong correlation between parameters *η* and *J* is inherent and not influenced by the inference procedure or model assumptions. This is in line with previous findings that have reported a strong and robust correlation between firing rates and synaptic weights across different brain states, environments and situations (Buzsáki and Mizuseki, 2014).

Using high-performance computing, model simulations can be run independently, creating a large training dataset for training deep neural density estimators in SBI approach (Hashemi et al., 2023). In contrast, HMC is limited to embarrassingly parallel execution with only independent chains on computational nodes (Hashemi et al., 2021). Moreover, when dealing with bistability in the state-space representation, HMC methods require significant computational time to detect state transitions in the latent space (see Table 1, Table 2, and Table 3) or need to be augmented with generative models such as normalizing flows (Hoffman et al., 2019; Gabrié et al., 2022). On the other hand, SBI offers efficient Bayesian estimation, even without detailed knowledge of the system’s state-space representation. This aligns with findings from recent studies that highlight the efficiency of SBI across various challenging inverse problems (Gonçalves et al., 2020; Deistler et al., 2022; Boelts et al., 2022, 2023; Hashemi et al., 2023; Lavanga et al., 2023; Yalccinkaya et al., 2023; Rabuffo et al., 2023; Sorrentino et al., 2023). By benchmarking and addressing questions such as computational cost, uncertainty quantification, inter-dependency exploration, and data availability, we conclude that SBI is more efficient than alternatives in making informed choices from microscopic states to emergent dynamics at the macro scale.

One of the main findings of this study is the effectiveness of deep neural networks in generating probability distributions for parameters of networks of spiking neurons. This approach outperforms other computational algorithms, such as MCMC, particularly in real-world applications involving missing data in bistable systems (see Figure 4). The effectiveness of deep generative models for inference from the mechanistic model of networks of spiking neurons is confirmed by the robustness of the estimation under significant dynamic noise (Figure 3, and Figure S7), as well as the precise estimation of input current (Figure 7) and the consistency with linear stability analysis (Figure 8). However, it is important to acknowledge that the use of deep generative models can result in an overestimation of uncertainty and correlations between parameters. To address this challenge, incorporating time-delay embedding is an effective remedy (Figure 5).

This study highlights the use of deep Neural ODEs in inferring vector fields at a macroscopic level, enabling the prediction of system dynamics from microscopic states. This approach has the potential to make interpretable predictions at larger scales from simulations at a detailed level, aiding in the prognosis and diagnosis of brain diseases. Recently, Sip et al. (2023) introduced a method using variational autoencoders for nonlinear dynamical system identification at the whole-brain level to infer both the neural mass model and the region- and subject-specific parameters from the functional data while respecting the known network structure. The scalability of Neural ODEs at the whole-brain network level remains to be investigated in future studies. In addition, symbolic regression applied to the outcomes from Neural ODEs may unveil closed-form equations for neural mass models, offering promising avenues for future research. This approach may lead to the discovery of concise data-driven mean-field representations of complex neural dynamics, contributing to our understanding of brain function.

In conclusion, this work highlights the improved accuracy and efficiency that deep learning techniques bring to the inference from networks of spiking neurons. It opens up exciting possibilities for future research in neural computation, where the trade-off between accuracy and uncertainty needs to be carefully considered. As we continue to explore the capabilities of SBI and Neural ODEs at larger scales, this study serves as a valuable step forward in our quest to unravel the complexities of neural networks and their computational mechanisms.

## Information Sharing Statement

All code is available at GitHub (https://github.com/ins-amu/Inference_MFM).

## Abbreviations

MF: Mean-Field
QIF: Quadratic Integrate-and-Fire
SBI: Simulation-Based Inference
SNPE: Sequential Neural Posterior Estimation
MCMC: Markov Chain Monte Carlo
HMC: Hamiltonian Monte Carlo
MAP: Maximum a Posteriori
DE: Differential Evolution
PSO: Particle Swarm Optimization
BO: Bayesian Optimization
ODEs: Ordinary Differential Equations
SDEs: Stochastic Differential Equations
Neural ODEs: Neural Ordinary Differential Equations

## Acknowledgements

This research has received funding from EU’s Horizon 2020 Framework Programme for Research and Innovation under the Specific Grant Agreements No. 945539 (Human Brain Project SGA3), No. 101147319 (EBRAINS 2.0 Project), and No. 101137289 (Virtual Brain Twin Project). The funders had no role in study design, data collection and analysis, decision to publish, or preparation of the manuscript. We thank Lionel Kusch and Spase Petkoski for fruitful discussions.

## Author contributions

Conceptualization: M.W., V.J., and M.H. Methodology: M.W., and M.H. Software: M.W., and M.H. Investigation: N.B., M.B., and M.H. Visualization: N.B., M.B., and M.H. Supervision: M.H., ad V.J. Funding acquisition: V.J. Writing - original draft: N.B., and M.H. Writing - review & editing: N.B, M.B, M.W., V.J, and M.H.

## 5. Supplementary

**Figure S1.**
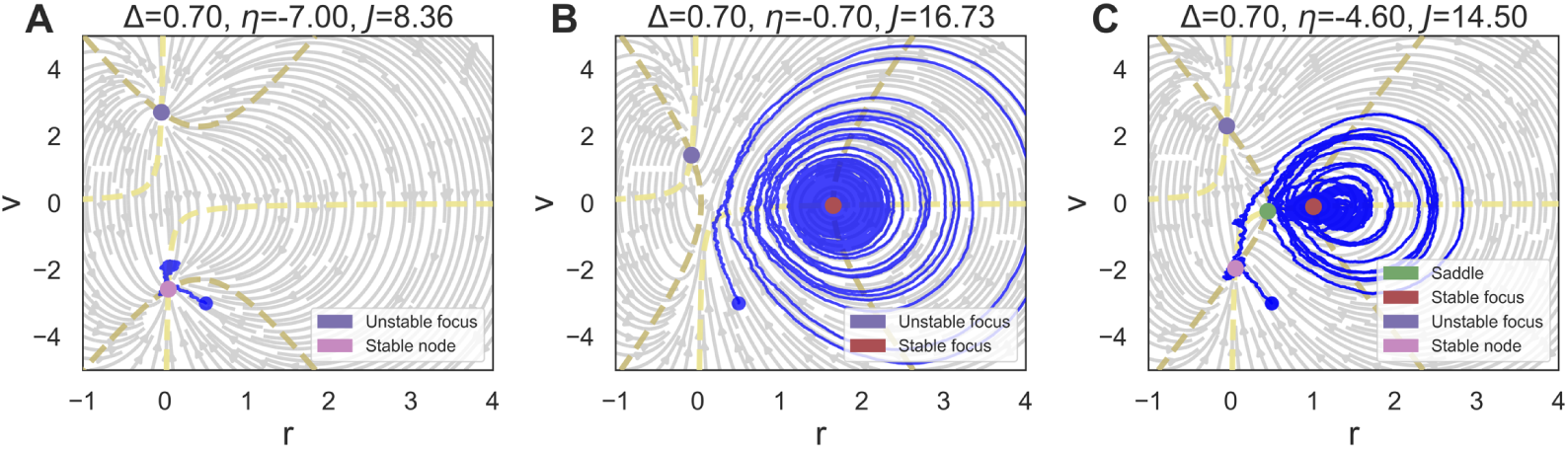
The phase-plane analysis of a mechanistic model of a network of all-to-all connected QIF neurons, using linear stability analysis. The stability of the system depends on the mean excitability (*η*) and synaptic weight (*J*), both normalized by the width of the input distribution (Δ). (**A**) A single stable node corresponding to a low-activity state. (**B**) A single stable focus (spiral) corresponding to a high-activity state. (**C**) A bistability between low and high firing rates (stable node and stable focus, respectively). The upper section between *v*- and *r*-nulclines (in dark and light yellow, respectively) corresponds to an unstable focus.

**Figure S2.**
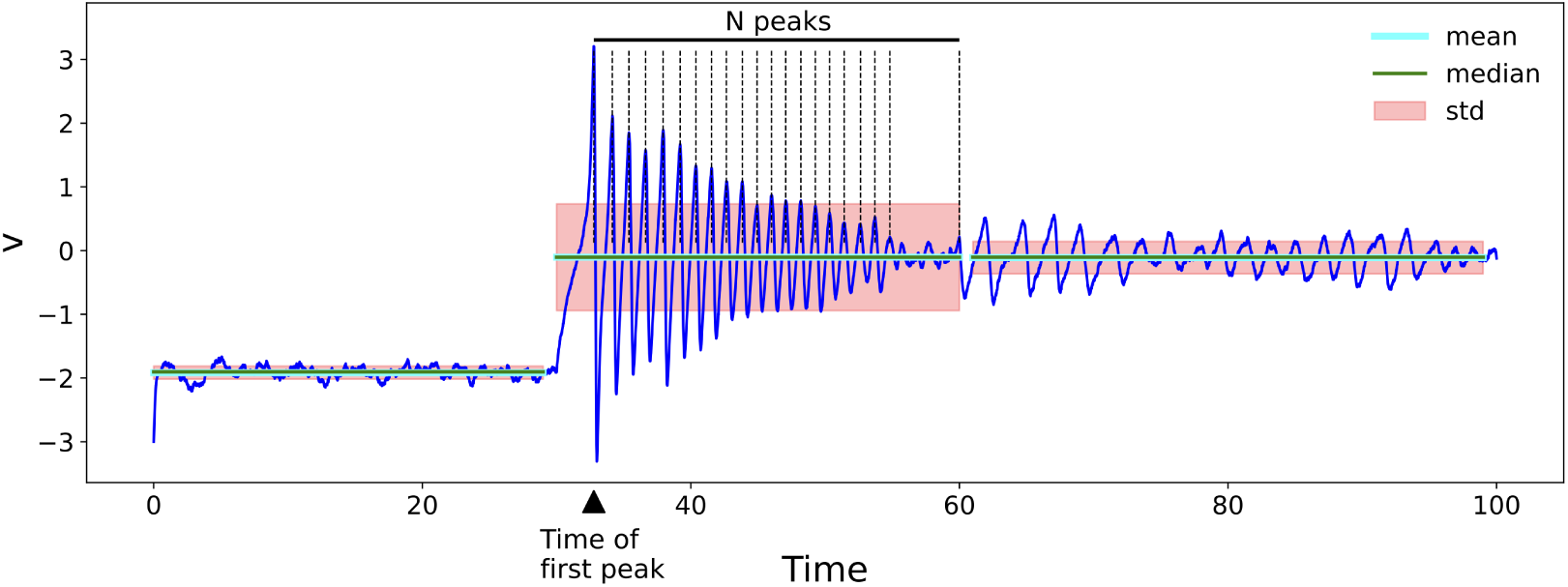
Data features extracted from mean membrane potential data. Additional data features that are not represented on this diagram include skewness and kurtosis.

**Figure S3.**
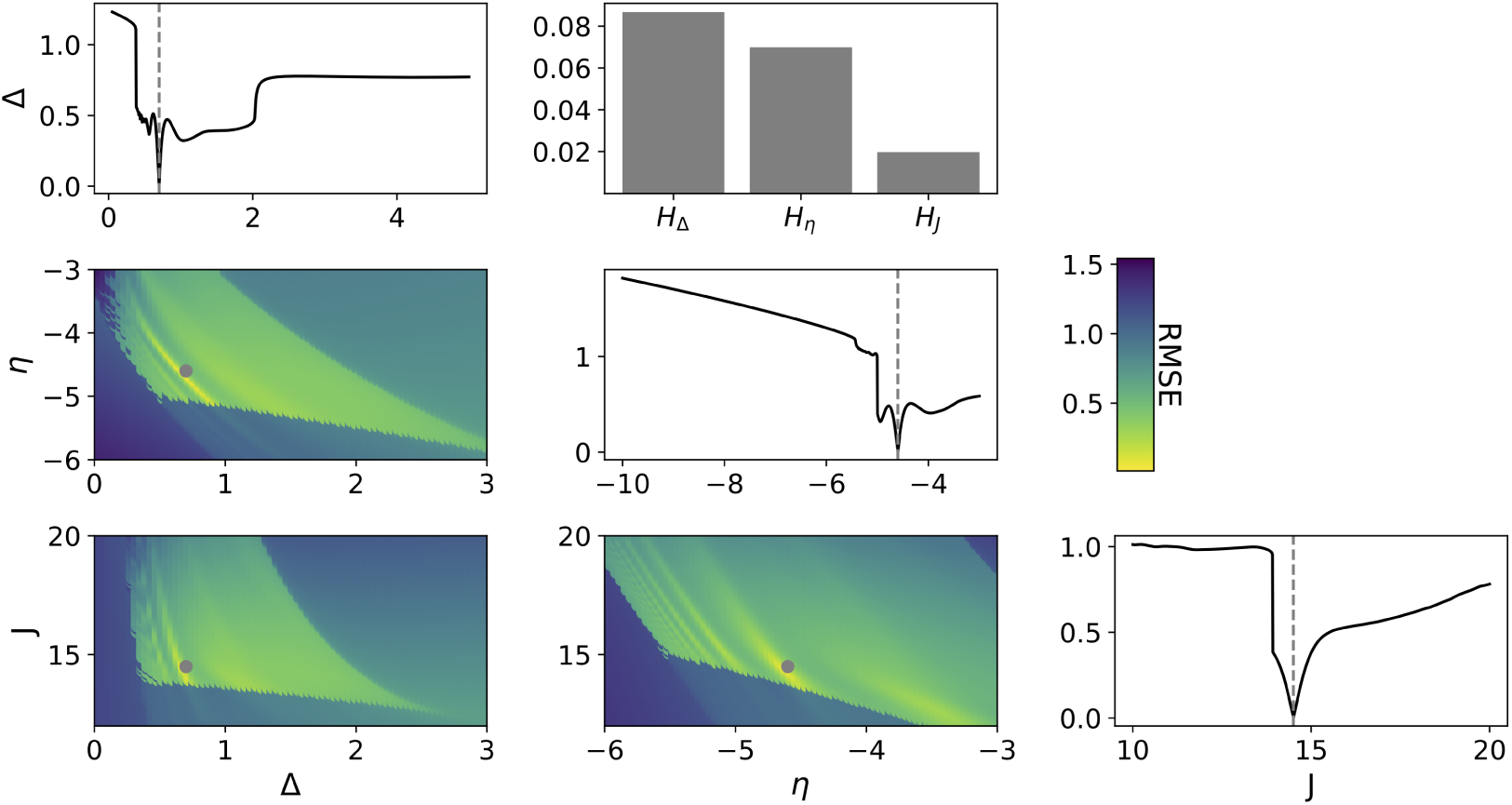
Sensitivity analysis on model parameters. The profile likelihood is calculated using the root-mean-squared error (RMSE) between the observed and generated data, considering either a single parameter (represented by black curves) or multiple parameters to vary (represented by colored surfaces), while keeping the other parameters fixed. The true parameters are shown in gray, represented by a dashed vertical line (for a single parameter) or 2D coordinates (for multiple parameters). The value of the Hessian matrix at the global minimum is displayed in the bar plot.

**Figure S4.**
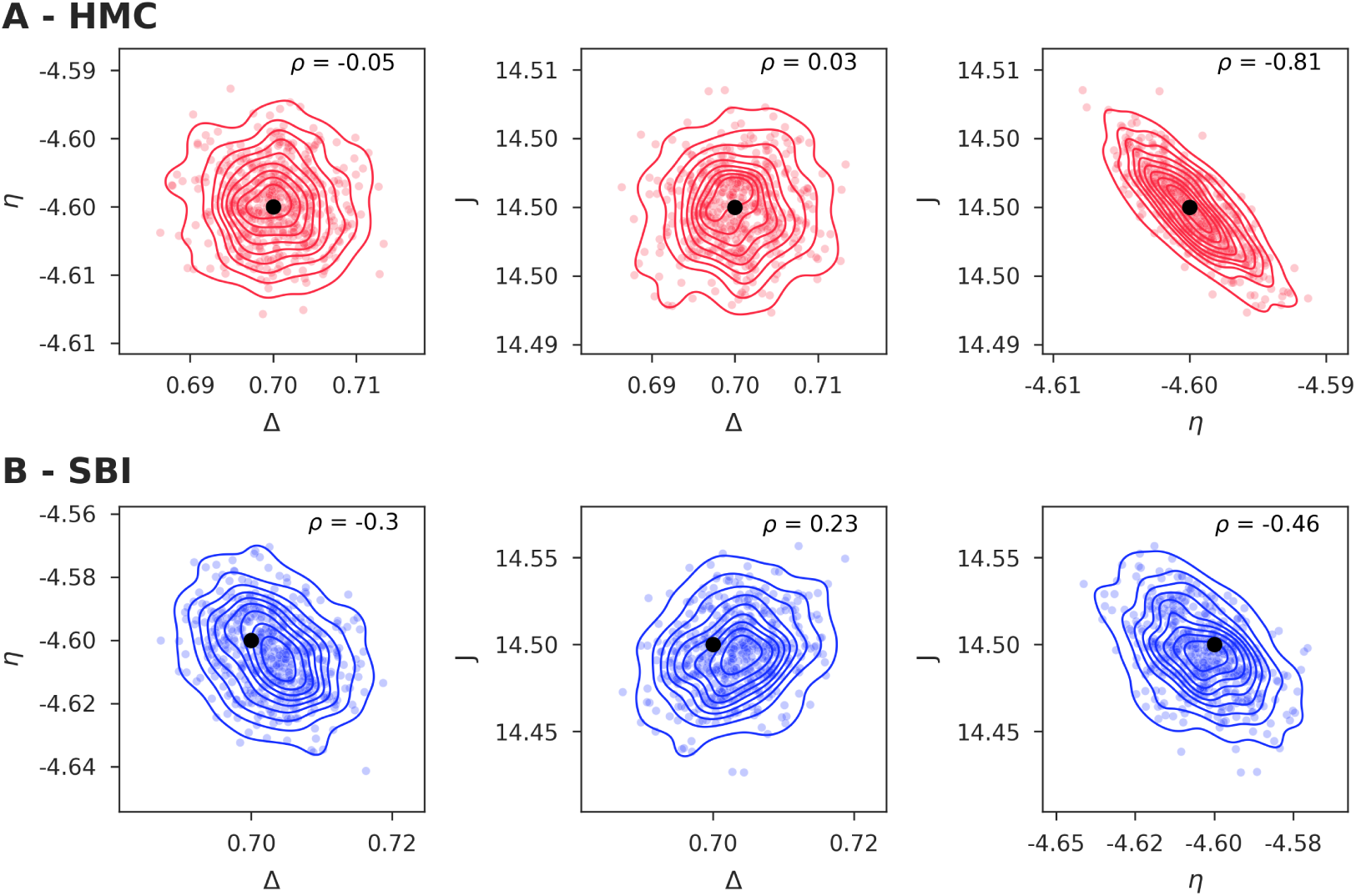
Paired posterior samples of model parameters inferred from deterministic data, using (**A**) HMC and (**B**) SBI. Samples do not exhibit significant pair-wise correlation in parameter couples (Δ, *η*) and (Δ, *J*). HMC manifests high linear correlation in sampling from joint parameters (*η*, *J*). In terms of uncertainty quantification, HMC offers a more informative posterior distribution compared to SBI. In terms of computational cost, SBI is approximately 60 orders of magnitude faster than HMC.

**Figure S5.**
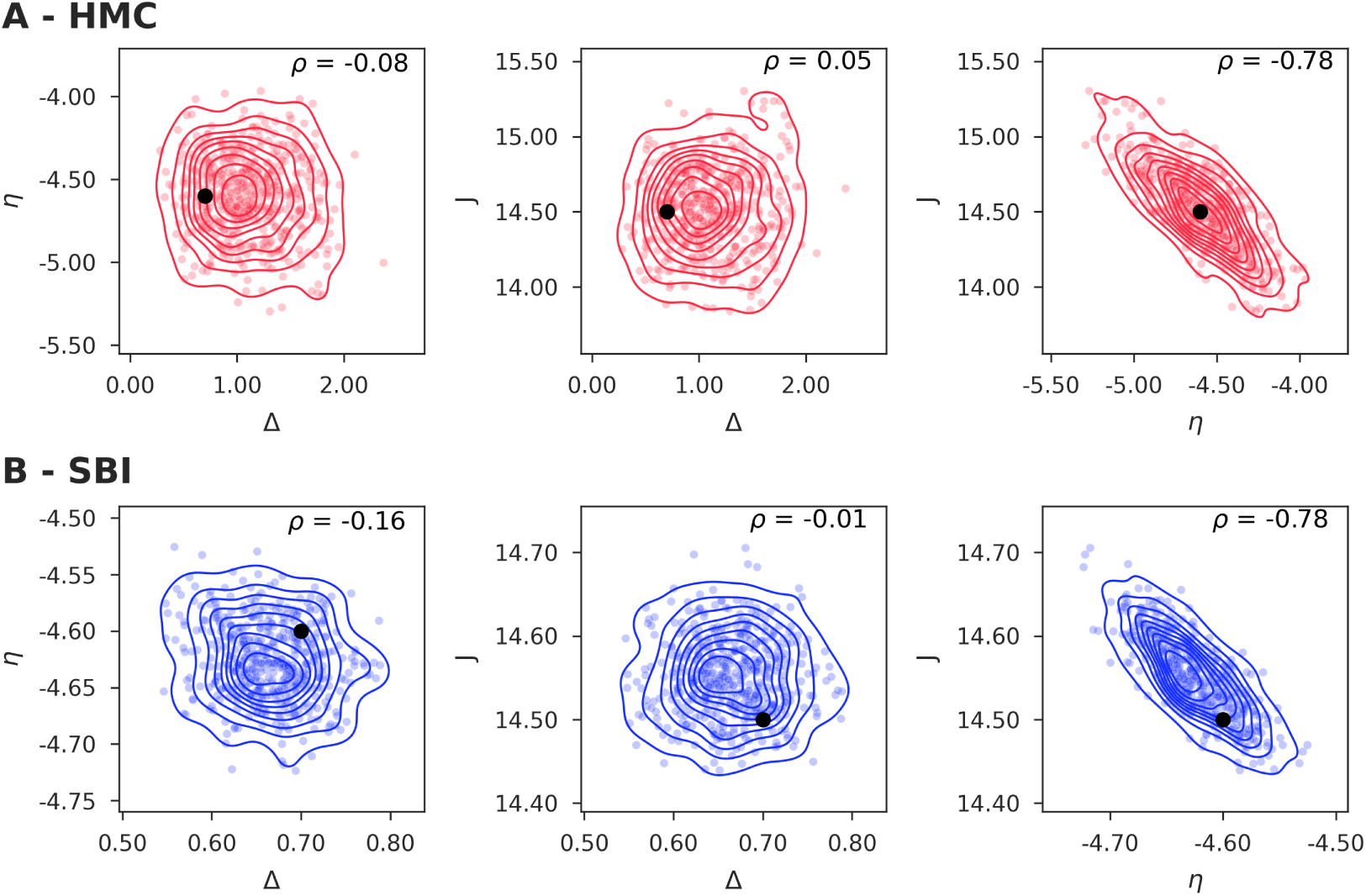
Paired posterior samples of model parameters inferred from noisy data, using (**A**) HMC and (**B**) SBI. Both algorithms provide uncorrelated samples for parameter couples (Δ, *η*) and (Δ, *J*) but exhibit the same level of strong linear correlation in sampling from joint parameters (*η*, *J*). Compared to HMC, SBI generates posteriors that more tightly center around the ground-truth parameters. Moreover, SBI remains approximately 8 orders of magnitude faster than HMC.

**Figure S6.**
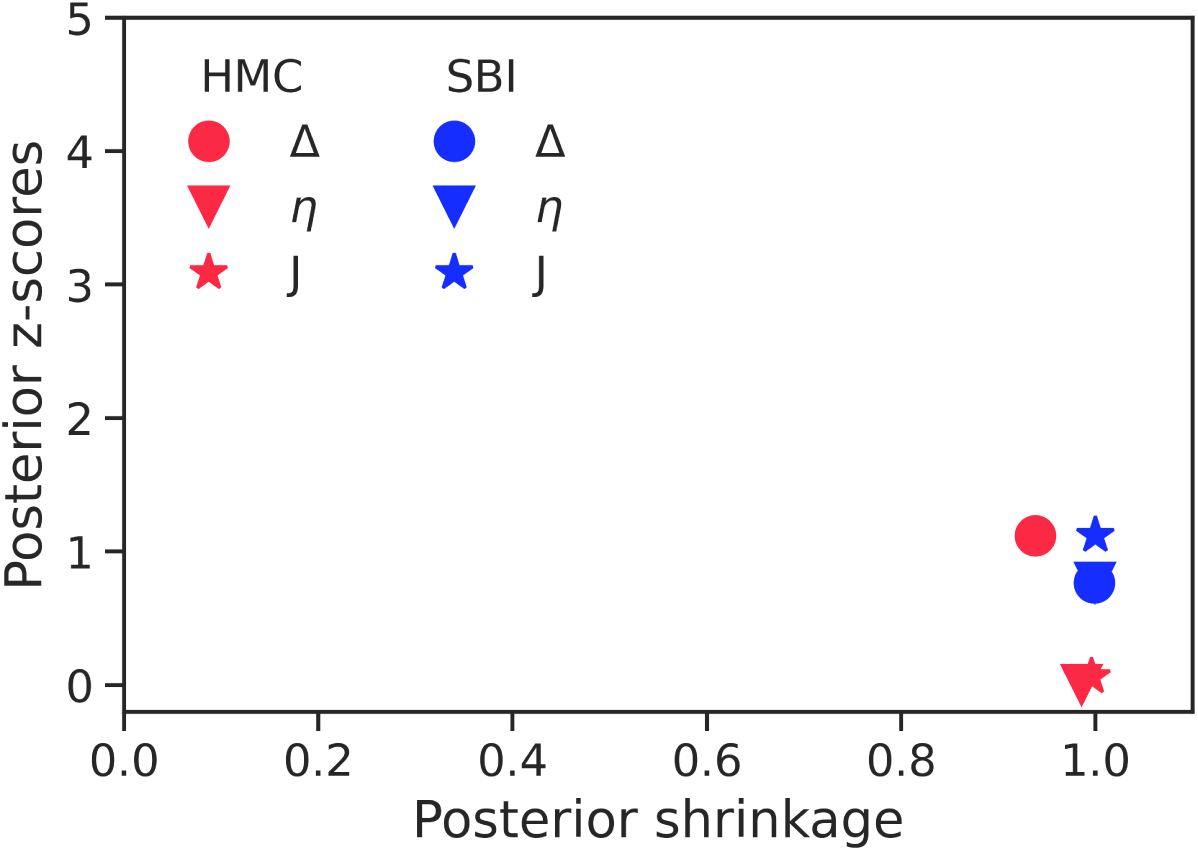
Posterior z-scores versus shrinkage, as a diagnostic for the reliability of Bayesian inference. An ideal Bayesian inference yields small z-scores (indicating less error) and high posterior shrinkage (indicating more contraction with respect to the prior distribution after learning from data). Therefore, their concentration lies in the right-bottom corner of the plot. When no data is missing, both HMC (red) and SBI (blue) perform near-optimally. However, SBI is significantly faster than HMC (at least 8 orders of magnitude). The posterior z-scores is defined as 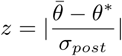, where *θ̄* and *θ^*^* are the posterior mean and the true values, respectively, whereas *σ_prior_*, and *σ_post_* indicate the standard deviations of the prior and the posterior, respectively. The posterior shrinkage defined as 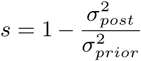, where *σ_prior_*, and *σ_post_* indicate the standard deviations of the prior and the posterior, respectively.

**Figure S7.**
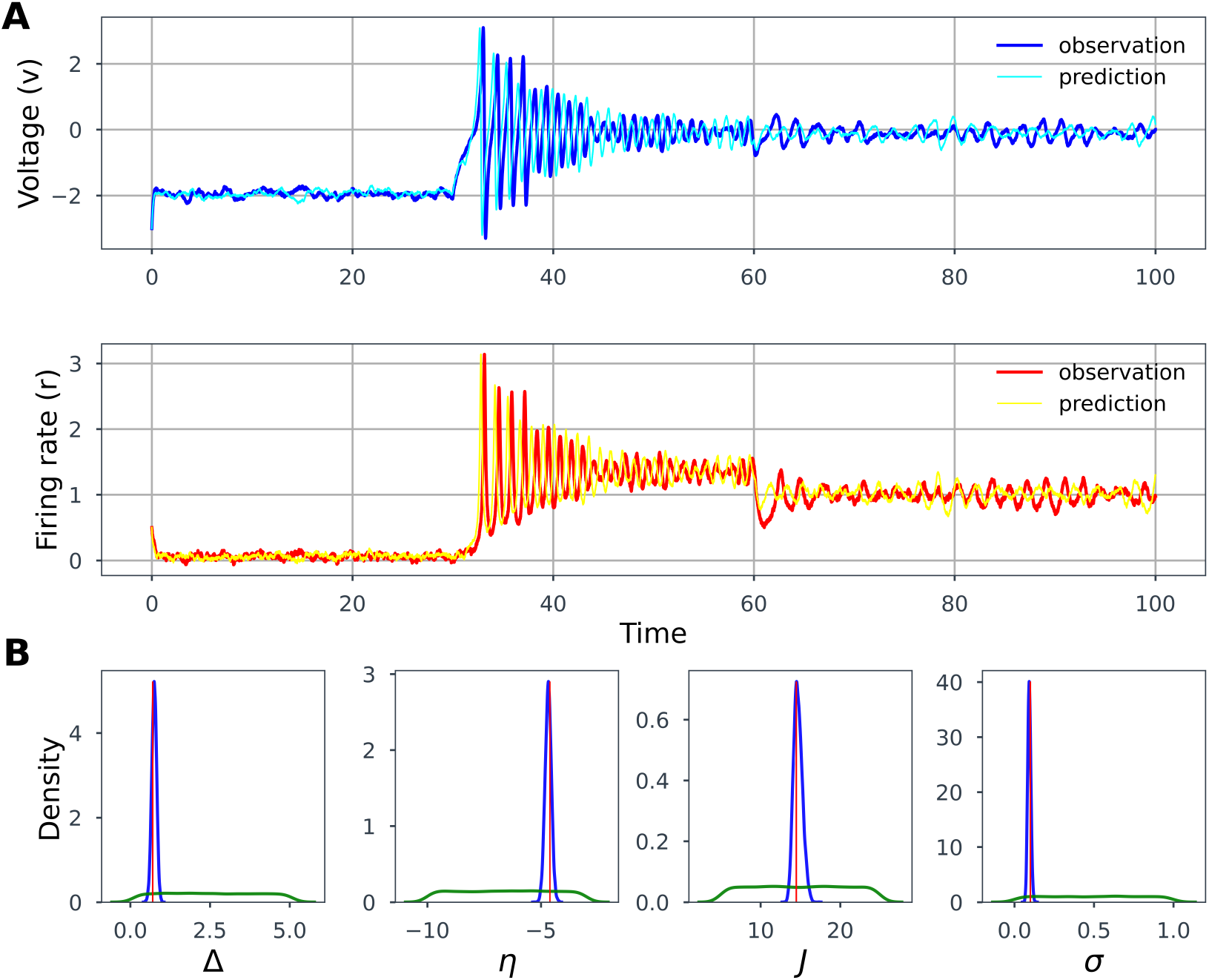
SBI over noisy data. (**A**) The time-series prediction of membrane potential (*v*) and firing rate (*r*). (**B**) Estimated posteriors of parameters Δ, *η*, *J*, and the intensity of dynamical noise *σ*. The true values are shown by vertical red lines. The priors and estimated posteriors using SBI are shown in green and blue colors, respectively. Given the low-dimensional data features of time-series, SBI is able to accurately and efficiently estimate the MF parameters including the dynamical noise in the system.

**Figure S8.**
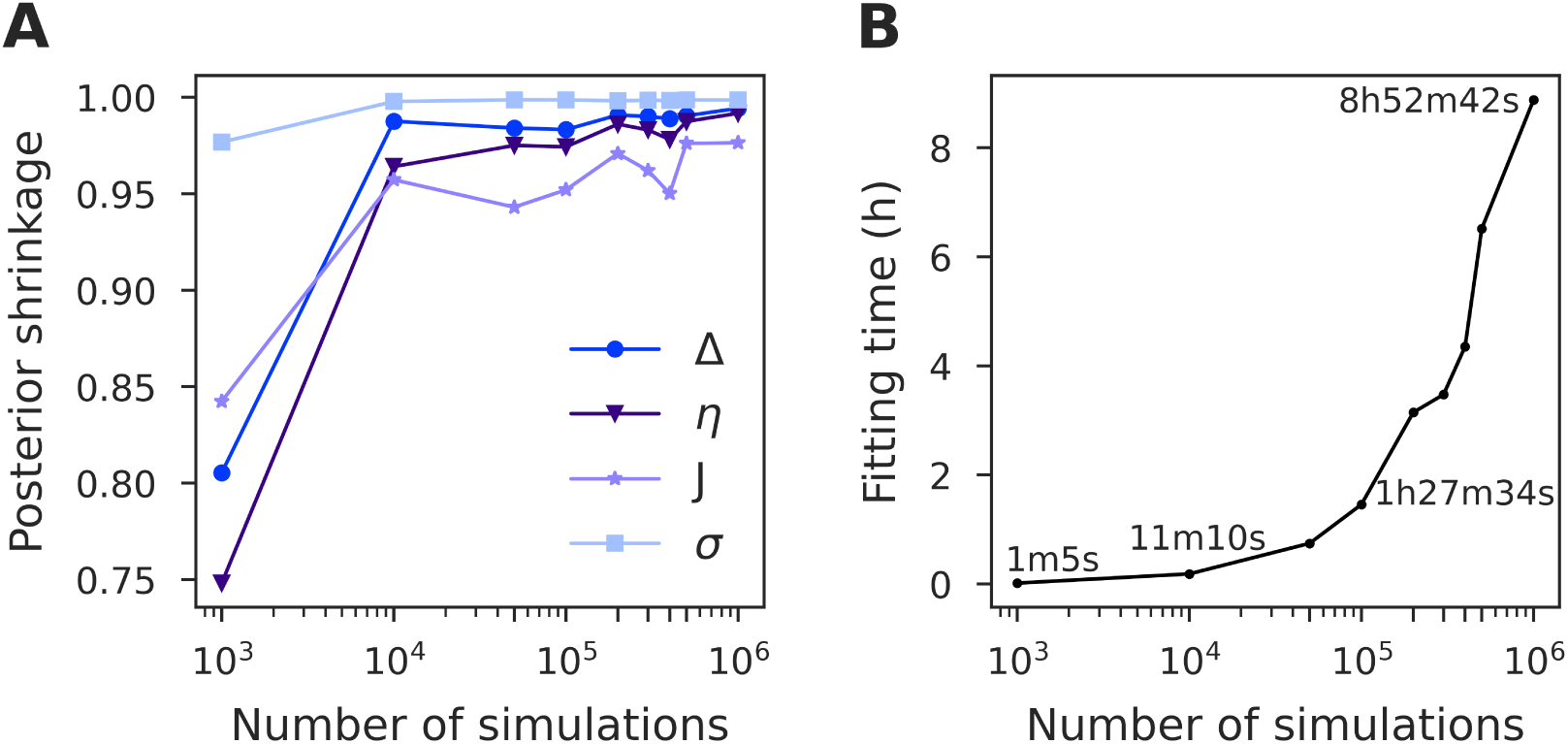
Performance of SBI as a function of the number of simulations for the training step. (**A**) The tendency of shrinkage towards one indicates that all the posteriors are well-identified. While the shrinkage of the posterior is significantly improved when increasing the number of simulations from 1k to 10k, further increasing it beyond 100k only results in a marginal improvement. (**B**) The computational cost for SBI, which includes the simulation, training, and sampling steps, increases exponentially with respect to the number of simulations.

**Figure S9.**
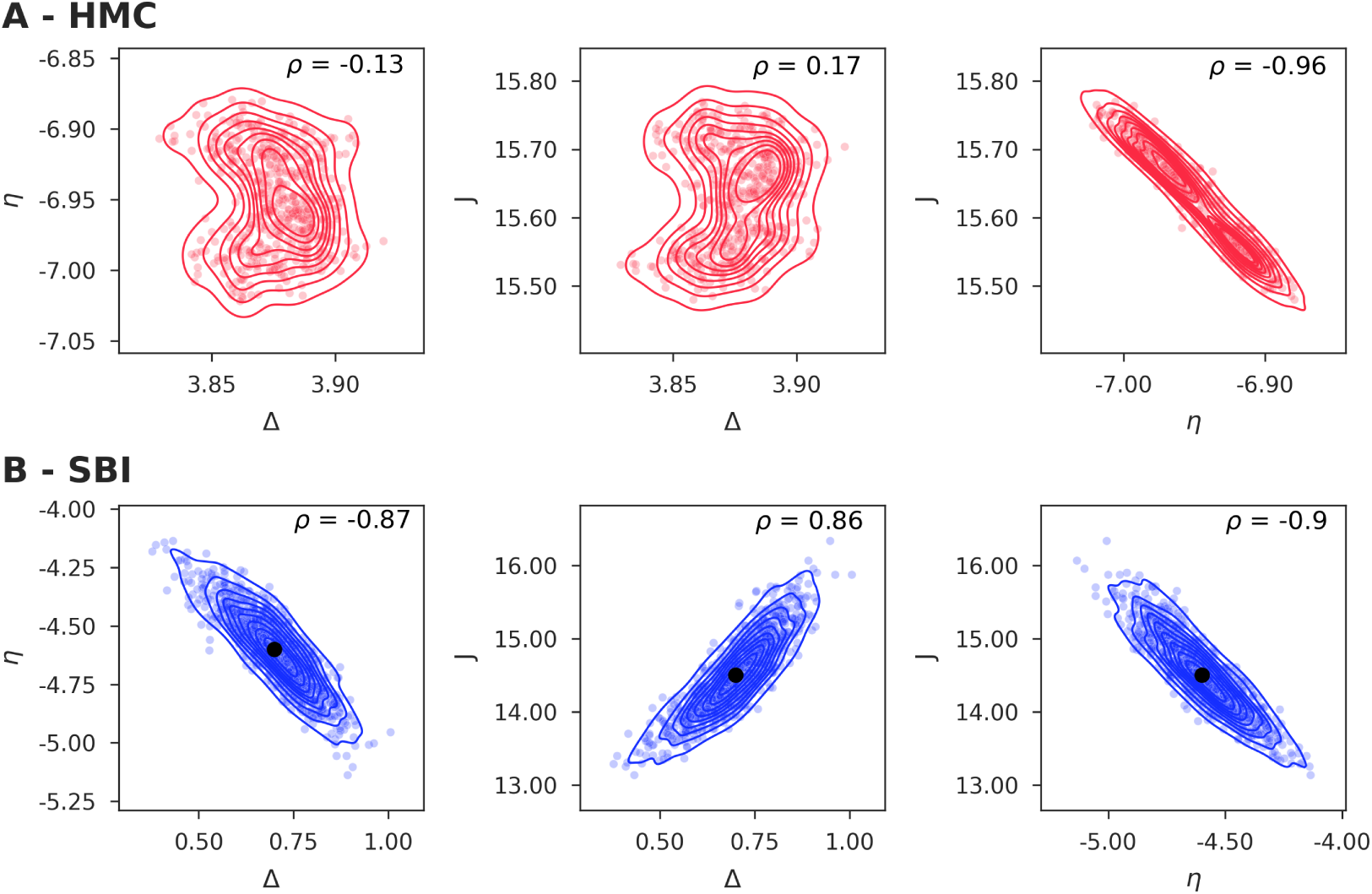
Paired posterior samples of model parameters inferred from noisy observation with missing data, using (**A**) HMC and (**B**) SBI, when only one variable (mean membrane potential *v*) is available. HMC samples deviate considerably from the true values, while SBI samples prove still reliable under such conditions, although accompanied with high correlation. Interestingly, SBI is approximately 68 orders of magnitude faster than HMC.

**Figure S10.**
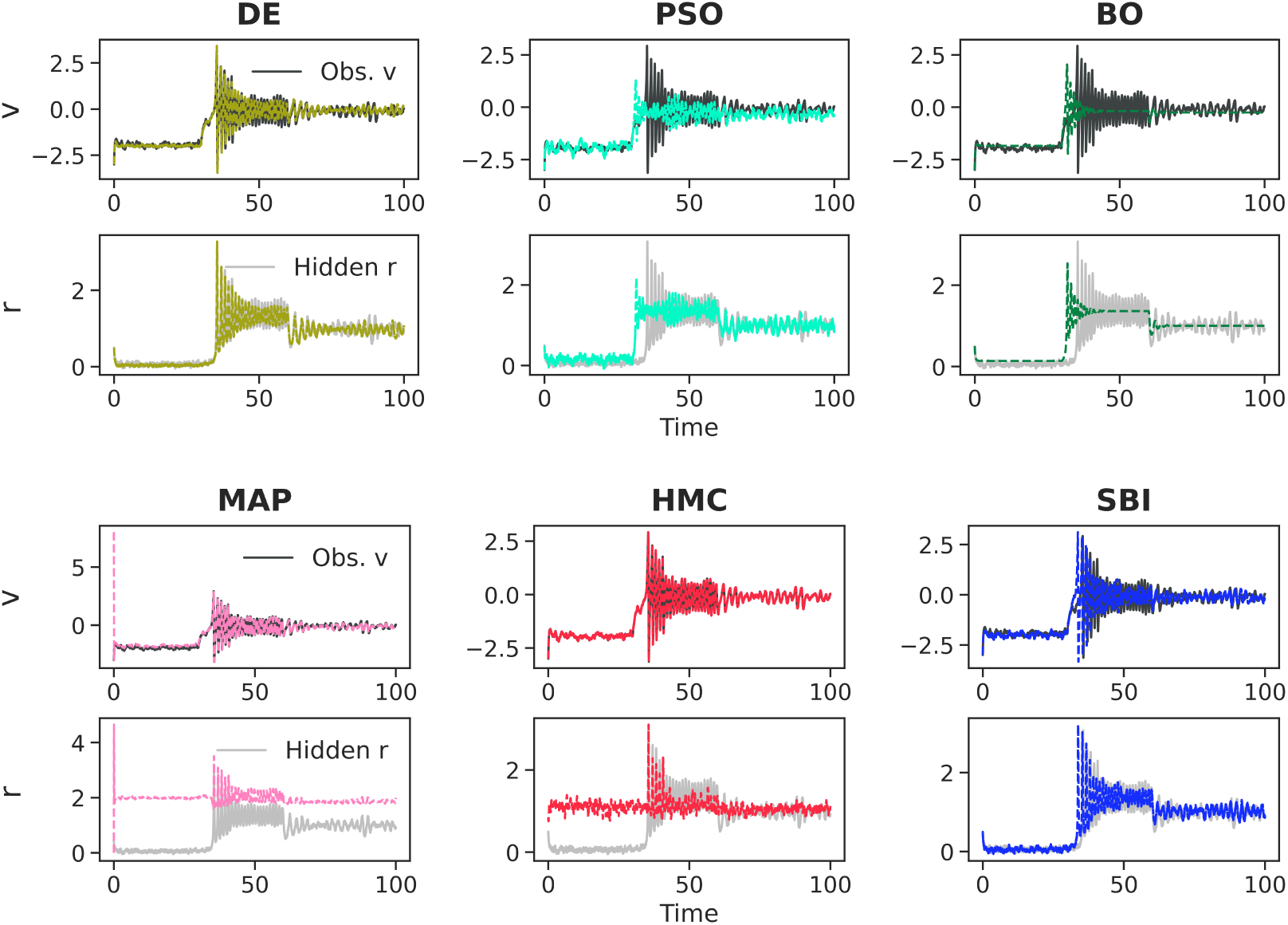
Fit to the noisy observation with missing data, i.e., when only one variable (mean membrane potential *v*) is available: observation *v* (top, black) and hidden *r* (grey, bottom). The time-series fit provided by the different inference algorithms is plotted in colors.

**Figure S11.**
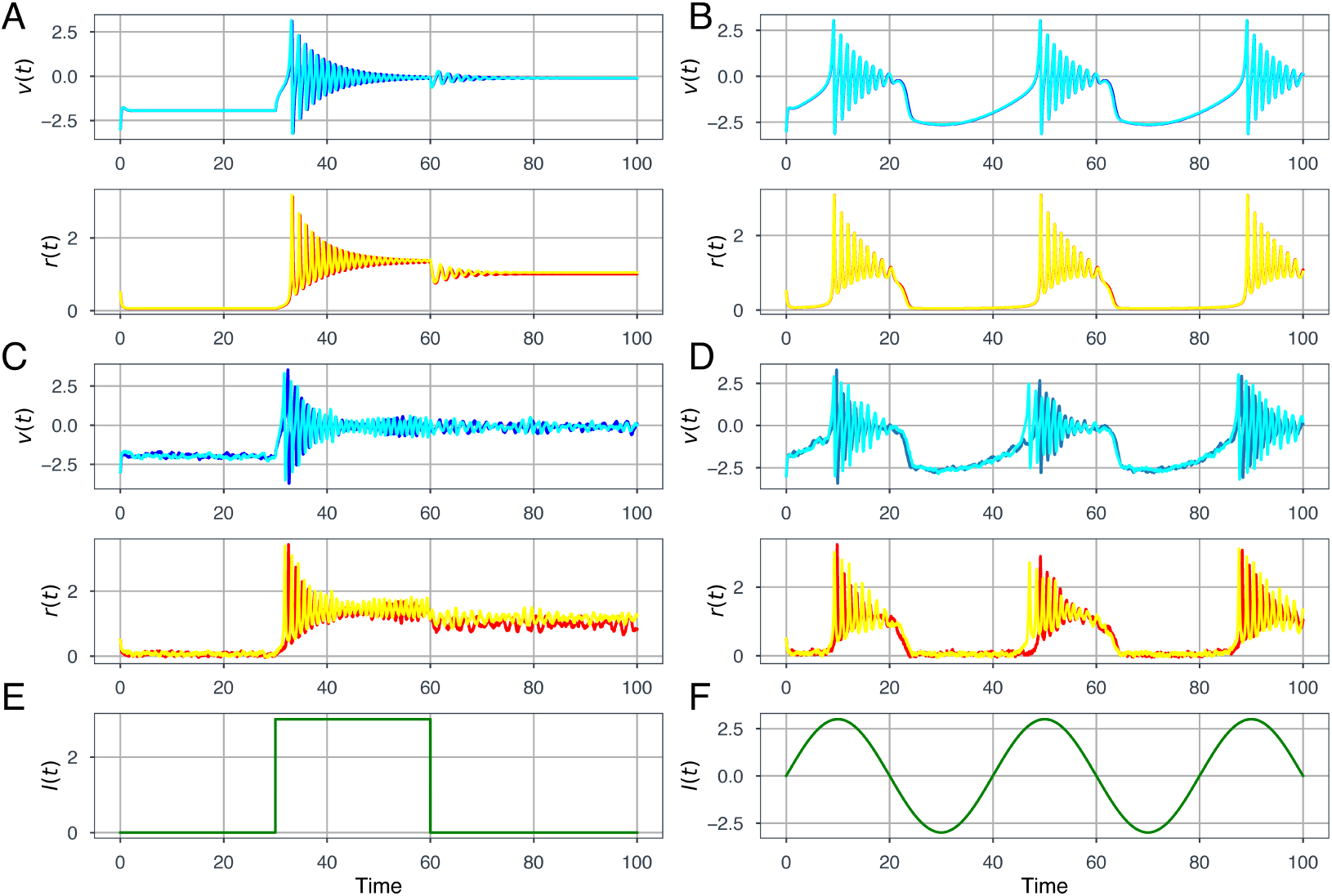
Exemplifying observed (*v* in blue and *r* in red) and predicted (*v* in cyan and *r* in yellow) time-series using SBI on (**A**, **B**) Deterministic data, (**C**, **D**) Stochastic data, given the input as (**E**, **F**) Step current with *I*(*t*) = *I*_0_ *·***1**stim(*t*), and sinusoidal current with *I*(*t*) = sin(*ω*_0_*t*)*·***1**stim(*t*), respectively.

**Figure S12.**
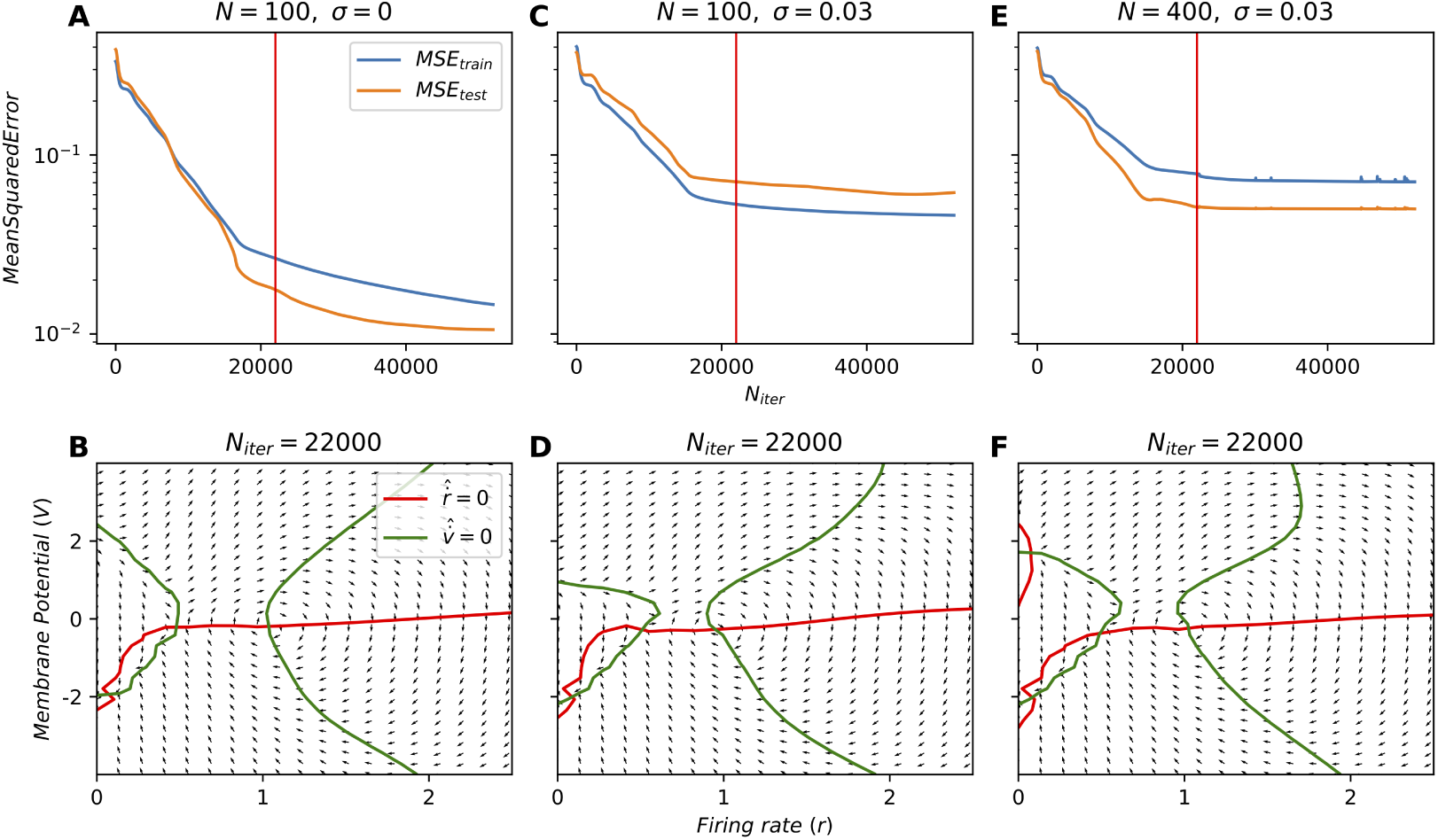
Loss functions for the training and test data using Neural ODEs on MF model. (**A**, **B**) For the deterministic training set of size 100, the phase-space estimation is performed at the 22000th iteration. (**C**, **D**) Training set of size 100 with dynamical noise. (**E**, **F**) Training set of size 400 with dynamical noise. The blue line represents the training error, while the orange line represents the test error. The vertical red line indicates the iteration at which a snapshot of the training is provided in the plot below.

**Figure S13.**
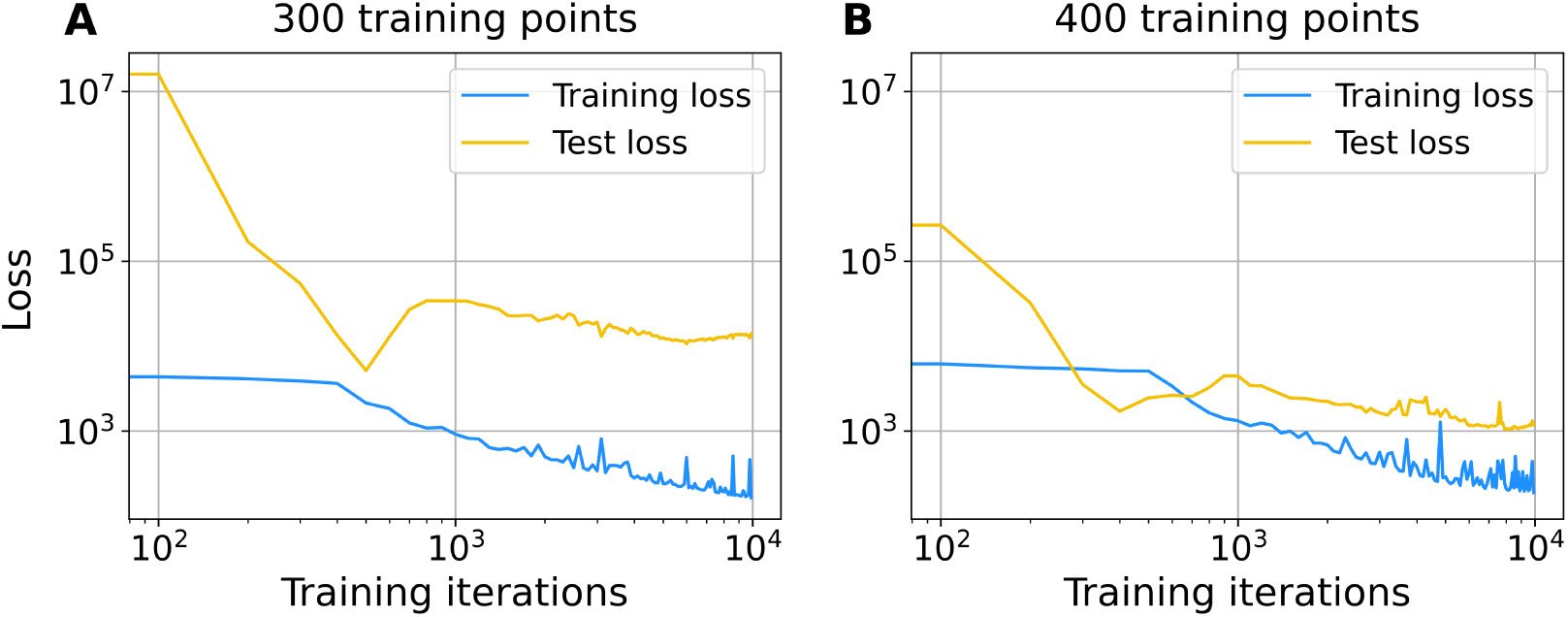
The loss function by training Neural ODEs on QIF data. The train and test quadratic loss function is calculated, when the train set is made of (**A**) 300 first points and (**B**) 400 first points. In each case, 10, 000 iterations were used for training the model.

## Notes

### Competing Interest Statement

The authors have declared no competing interest.

